# Invertebrate-induced priority effects and secondary metabolites control fungal community composition in dead wood

**DOI:** 10.1101/2021.12.01.470726

**Authors:** Lisa Fagerli Lunde, Rannveig Jacobsen, Håvard Kauserud, Lynne Boddy, Line Nybakken, Anne Sverdrup-Thygeson, Tone Birkemoe

## Abstract

During decomposition of organic matter, microbial communities may follow different successional trajectories depending on the initial environment and colonizers. The timing and order of the assembly history can lead to divergent communities through priority effects. We explored how assembly history and resource quality affected fungal dead wood communities and decomposition, 1.5 and 4.5 years after tree felling. Additionally, we investigated the effect of invertebrate exclusion during the first two summers. For aspen (*Populus tremula*) logs, we measured initial resource quality of bark and wood, and surveyed the fungal communities by DNA metabarcoding at different time points during succession. We found that a gradient in fungal community composition was related to resource quality and discuss how this may reflect tolerance-dominance trade-offs in fungal life history strategies. As with previous studies, the initial amount of bark tannins was negatively correlated with wood decomposition rate over 4.5 years. The initial fungal community explained variation in community composition after 1.5, but not 4.5 years, of succession. Although the assembly history of initial colonizers may cause alternate trajectories in successional communities, our results indicate that the communities may easily converge with the arrival of secondary colonizers. We also identified a strong invertebrate-induced priority effect of fungal communities, even after 4.5 years of succession, thereby adding crucial knowledge to the importance of invertebrates in affecting fungal community development. By measuring and manipulating aspects of assembly history and resource quality that have rarely been studied, we expand our understanding of the complexity of fungal community dynamics.

## Introduction

Biological communities developing in newly emerged habitats depend on the interplay between the biotic and abiotic environment. The community assembly can lead to stable or alternate community compositions, that is, they either converge or diverge over time (Chase, 2010; Fukami, 2015; Young, Chase, & Huddleston, 2001). This *historical contingency*, or unforeseen outcomes from past events, is inherently driven by deterministic and stochastic effects of the initial environment (Ben-Menahem, 1997). Perhaps the most well known example of a stochastic effect is assembly history, i.e. the initial timing of species colonizations, which ultimately can determine the subsequent community structure in processes called *priority effects* (Chase, 2003; Drake, 1991; Fukami, 2015). An example of deterministic effects is when environmental conditions or biotic interactions exclude species from the community after they have arrived, although the two are not independent of one another (Cadotte & Tucker, 2017; Kraft et al., 2015; Vellend, 2010). Understanding the processes that govern community assembly is important for detecting drivers of biological communities which ultimately link to ecosystem functions.

Dead wood provides diverse habitats which are important for global biodiversity and carbon sequestration (Goodale et al., 2002; Seibold et al., 2021; Stokland, Siitonen, & Jonsson, 2012). The organisms that are dependent on dead wood comprise mostly fungi and invertebrates – particularly insects. Fungi are by far the most effective decomposers of dead wood through their lignocellulolytic abilities (Baldrian, 2006; Rayner & Boddy, 1988). Although wood decomposition is a process that sometimes lasts for decades (Stokland et al., 2012), most experimental studies that try to assess the importance of dead wood communities have focused only on the first few years of decomposition (e.g. Seibold et al., 2021; Ulyshen, 2016; Venugopal, Junninen, Edman, & Kouki, 2017). However, given the central role that fungi have in nutrient cycling in forests, it is crucial to expand this successional perspective.

Fungal community development in dead wood is governed by environmental conditions (e.g. water content, temperature) and biotic interactions, particularly combative interactions between fungi (Lynne Boddy, 2000; Lynne Boddy & Heilmann-Clausen, 2008; Rayner & Boddy, 1988; Venugopal et al., 2017). The importance of environment is reflected by the different life history strategies that dead wood fungi have evolved: stress-tolerant, ruderal and competitive (Lynne Boddy, 2001; Lynne Boddy & Heilmann-Clausen, 2008; Treseder & Lennon, 2015). This is exemplified by the distinct fungal communities that are hosted by different tree species (e.g. Lynne Boddy & Heilmann-Clausen, 2008; Norberg, Halme, Kotiaho, Toivanen, & Ovaskainen, 2019) which probably reflects inherent physicochemical qualities of the wood (Hoppe et al., 2016), including pH, nutrient and secondary metabolite content, and the selective advantages of fungal life history strategies (Rayner & Boddy, 1988).

The amount and type of inhibitory secondary metabolites which the tree produces give a competitive advantage to fungal species that tolerate this stress (Valette, Perrot, Sormani, Gelhaye, & Morel-Rouhier, 2017). Among the most common secondary metabolites are phenolic compounds, like tannins and flavonoids, which inhibit growth of many species of fungi (Flores & Hubbes, 1979; Hammerbacher, Raguschke, Wright, & Gershenzon, 2018) and the rate of decomposition (Coq, Souquet, Meudec, Cheynier, & Hättenschwiler, 2010; Loranger, Ponge, Imbert, & Lavelle, 2002; Valette et al., 2017). The concentrations of phenolic compounds can differ markedly between and within individual trees of the same species (T Birkemoe et al., 2021). However, few studies (Bailey et al., 2005; T Birkemoe et al., 2021) have investigated whether such intraspecific variation in secondary metabolite concentration can govern community assembly of fungi in wood.

The initial fungal colonizers of wood arrive rapidly as spores or are latently present in functional sapwood (Lynne Boddy & Hiscox, 2016; L Boddy, Hiscox, Gilmartin, Johnston, & Heilmann-Clausen, 2017). A wide diversity of fungi, including wood decaying species, are present as latent propagules, but relatively few of them establish overtly when the sapwood becomes dysfunctional (Chapela & Boddy, 1988b; Cline, Schilling, Menke, Groenhof, & Kennedy, 2018; Hiscox, Savoury, Müller, et al., 2015; Parfitt, Hunt, Dockrell, Rogers, & Boddy, 2010). These initial colonizers of wood are, however, soon replaced by more combative fungi arriving as spores or mycelial cords (Chapela & Boddy, 1988a; Hiscox, O’leary, & Boddy, 2018). The outcome of such competitive interactions can be altered by invertebrate grazing (A’Bear, Jones, & Boddy, 2014; Crowther, Boddy, & Jones, 2011). Further, invertebrates can affect fungi in other ways (Tone Birkemoe, Jacobsen, Sverdrup-Thygeson, & Biedermann, 2018), especially by aiding in dispersal (e.g. bark beetles: Harrington, Furniss, & Shaw, 1981; Pettey & Shaw, 1986).

There is accumulating evidence that the developing community in dead wood can be shaped by priority effects through altered assembly history (e.g. Dickie, Fukami, Wilkie, Allen, & Buchanan, 2012; Fukami et al., 2010). For example, on the forest floor, pre-colonization of wood by *Stereum hirsutum* lead to distinct fungal communities compared to when seven other species were introduced first (Hiscox, Savoury, Müller, et al., 2015). There are also indications that the presence of invertebrates can lead to priority effects in fungal communities of dead wood (Rannveig M Jacobsen, Sverdrup-Thygeson, Kauserud, Mundra, & Birkemoe, 2018; Leopold et al., 2017). However, the few long-term studies that exist have only traced a limited number of fungal species (Rannveig Margrete Jacobsen, Birkemoe, & Sverdrup-Thygeson, 2015; Weslien, Djupström, Schroeder, & Widenfalk, 2011), rather than looking at the effect on the whole community. Given that wood decomposition is a slow process and priority effects might attenuate over time (Dickie et al., 2012; Fukami, 2015; Norberg et al., 2019), there is a need for long-term studies investigating how assembly history affects fungal community composition and wood decomposition.

In this study, we explore how resource quality and assembly history shape fungal community development in dead wood of aspen (*Populus tremula* L.). In particular, we quantify the initial secondary metabolites, carbon and nitrogen content, and identify the initial fungal community composition of the dead wood habitat. Furthermore, we manipulate assembly history by excluding invertebrates for two summers in an in-field experiment. To uncouple the effects of deterministic and stochastic processes caused by spatial variation on fungal communities, we transferred aspen logs to 30 different locations in two landscapes. To cover temporal variation in the fungal communities, the logs were sampled twice – after 1.5 and 4.5 years (termed year 2 and 5). By applying DNA metabarcoding of the ITS2 region, we ask how does: (1) *resource quality* (initial secondary metabolite, carbon and nitrogen content), *and* (2) *assembly history* (initial fungal community and invertebrate exclusion) *affect fungal community composition, OTU richness and wood decomposition?* Answering these questions will reveal important drivers of biological communities and dead wood decomposition.

## Materials and methods

### Study area

Seventeen aspen (*Populus tremula* L.) trees were felled from the same stand in ås municipality in Norway (59°66’N 10°79’E, 92 m.a.s.l.) during March 2014. The trees were cut into 1 m long logs (diameter (D) = 20.5–36.4 cm, mean = 27.6 cm), which were transported to 15 sites in each of two boreal forest landscapes in south-east Norway: Losby forest holdings in Østmarka (59°87’N 10°97’E, 250–300 m.a.s.l.) and Løvenskiold-Vækerø in Nordmarka (60°08’N 10°58’E, 300–500 m.a.s.l.). Both forests are dominated by spruce (*Picea abies* (L.) H.Karst.), with pine (*Pinus sylvestris* L.), birch (*Betula pubescens* Ehrh.) and aspen as subdominants. All sites were in mature, closed-canopy managed forest, with a mean distance between sites of 120 m in Østmarka and 276 m in Nordmarka.

### Wood and bark samples for analysis of the initial fungal community and resource quality

After felling, 53 discs were cut from the fresh wood between every two or three logs (Figure S1). The discs were used for DNA analysis and chemical measurements to identify the initial fungal community and determine resource quality, respectively. Aseptically (ethanol-and-flame-sterilization), bark was removed with a knife, then wood samples for DNA analysis were taken by drilling 10 cm into the wood with a 12-mm drill bit. Each sample consisted of woodchips from drilling at two different positions along the circumference of the section. Wood samples were stored at -80^°^C prior to DNA extraction. Wood samples for analysis of resource quality were taken next to the drilling holes where samples for DNA analysis had been taken. Samples were taken by drilling, using the same method as above, but followed by drying the samples at 30^°^C overnight and subsequently storing them at -20^°^C. Bark samples for analysis of resource quality were taken 3 months after felling, from logs stemming from the same 17 aspen trees and distributed in the same two forest landscapes. Bark samples were taken with a knife that was sterilized between each sample, subsequently dried at 30^°^C and stored at -20^°^C. T Birkemoe et al. (2021) describes the sampling of these logs in detail.

### Invertebrate exclusion experiment

Four logs were placed at each of the 30 sites and subjected to the following treatments from April 2014 to November 2015: (1) Invertebrate exclusion (‘cage’) where logs were enclosed in a fine polyester plastic mesh (1×1 mm) net suspended from a frame. Colonization from below was prevented by a polyethylene plastic sheet beneath the logs. A mesh size of 1 mm effectively excludes macro- and larger meso-invertebrates, which we expect to have greatest impacts on fungi (Crowther, Boddy, et al., 2011; Crowther, Jones, & Boddy, 2011). Since the plastic sheets would prevent not only invertebrates, but also fungi from colonizing from the soil, these were included in all treatments. (2) ‘Cage control’ logs placed within cages identical to those in (1), except with four large holes (D = 20 cm) allowing invertebrate access. This treatment was to control for microclimatic effects of the cage itself. (3) Uncaged logs (‘control’) placed on a plastic sheet. (4) Uncaged ‘positive control’ logs baited with an ethanol bottle (1L, 96% ethanol) with small holes for evaporation to attract wood-inhabiting invertebrates. In November 2015, cages and plastic sheets were removed, but all logs were kept in the same positions on the ground. More information on the experimental set-up can be found in Rannveig M Jacobsen et al. (2018).

### Wood samples for analysis of the successional fungal communities

Wood samples for DNA analysis were taken from the logs at the end of the exclusion period in November 2015 (termed year 2), and again in November 2018 (termed year 5). The same methods as for fresh wood samples were used, but with drilling at three different positions along the circumference (top and both sides) for each sample. In year 2, each log was sampled 25 cm (end section) and 50 cm (mid section) from the end with the least damage to the bark. In year 5, samples were taken 20 cm (end section) and 45 cm (mid section) from the same log end.

### Wood samples for analysis of wood decomposition

Wood samples for density analysis were taken in November 2015 (year 2) and in May - June 2018 (year 5), next to the drilling holes for DNA analysis samples in the respective years. Unfortunately, we were not able to sample wood density and woodchips for DNA in parallel during 2018 as the risk of forest fire prevented flame sterilization during summer. Samples for density analysis were taken with a core sample drill (length = 10 cm, D = 12 mm) in two replicates – top and one side – for each section and pooled together for analysis. Only the outer 5 cm (without bark) of the core sample was used for analyses because the inner region was often too decayed to be sampled in year 5 (Tangnæs & Andelic, 2019). Wood density was calculated as dry mass divided by fresh volume. Fresh volume was first measured by water displacement, then the samples were oven-dried at 103^°^C overnight before measuring dry mass.

### Chemical analyses

The analyses of carbon, nitrogen and phenolic compounds are described in detail in T Birkemoe et al. (2021). Briefly, dried wood and bark samples were ground to powder with a mixer mill (Retsch MM400, Haan, Germany) from which subsamples for different analyses were taken. Carbon and nitrogen concentrations were analysed on a Micro Cube elemental analyser (Elementer Analysen, Hanau, Germany). Low-molecular phenolic compounds were extracted in methanol and subsequently analysed on HPLC (Agilent Series 1200, Agilent Technologies, Waldbronn, Germany). Compound identification was based on retention times, online UV spectra and co-chromatography of commercial standards (Nybakken, Hörkkä, & Julkunen-Tiitto, 2012). Individual phenolic compounds were assigned to chemical groups (phenolic acids, salicylates and flavonoids; T Birkemoe et al. (2021)). Two fractions of condensed tannins, methanol-soluble (MeOH-soluble) and methanol-insoluble (MeOH-insoluble), were quantified by the acid butanol assay for proanthocyanidins (Hagerman, 2002) from the liquid HPLC-extract and from the extraction residue, respectively.

### DNA isolation, amplification and sequencing

Samples in year 2 and immediately after felling (fresh wood; initial fungal community) were analysed for DNA as described in Rannveig M Jacobsen et al. (2018). Samples in year 5 were analysed for DNA in the same way, except for minor differences in centrifugation speed (always 14 000 x *g* in year 2 and fresh wood) and primer use (see below). In 50 mL Falcon tubes, 15 mL woodchips and 15 mL CTAB lysis buffer (pH 8.0; 100mM Tris-HCl, 1.4 M NaCl, 20 mM EDTA, 2% w/v CTAB) were added along with 7 stainless steel beads (5 mm). The tubes were homogenized in a mixer mill (Retsch MM400, Haan, Germany) at 30 Hz for 90 s, placed at -80°C for 30 min and then incubated at 54°C overnight. Next, samples were inverted, left to cool, and vortexed with 15 mL of chloroform. They were centrifuged for 25 min in a Rotina 46 (Hettich Zentrifugen, Tuttlingen, Germany) at 1780 x *g* until the phases had separated. In a clean tube, 5 mL of the upper phase was transferred along with 5 mL cold isopropanol. To precipitate the DNA, the tubes were inverted and placed at -20°C for 30 min. The tubes were centrifuged at 13 000 x *g* for 10 min in an Eppendorf 5804R (Eppendorf, Hamburg, Germany) and the supernatant was discarded. The remaining pellet was washed with 1000 µL cold 70% ethanol, vortexed, centrifuged for 3 min at 13 000 x g and discarded. Excess ethanol was decanted by 60°C incubation and dried on laboratory tissue paper. Finally, the DNA was resuspended with 60 µL milli-Q H2O. The extracts were cleaned using an E.Z.N.A.® Soil DNA kit (Omega Bio-tek, Norcross) as recommended by the manufacturers. Then, we washed again with 500 µL DNA Wash Buffer and eluted twice with 20 µL Elution Buffer, resulting in 40 µL suspended DNA.

Amplicon libraries were constructed from 252 DNA extracts, 14 technical replicates and three PCR negatives. They were prepared on three 96 PCR plates with uniquely tagged ITS2 primers in 20 µL volume samples: 6.2 µL milli-Q H2O, 4 µL 10x-dilution DNA template, 4 µL Phusion HF Buffer, 2 µL dNTPs (10 mM), 1 µL bovine serum albumin (BSA; 10 µg/µL), 0.6 µL dimethyl sulfoxide (DMSO), 0.2 µL Phusion Hot Start II High-Fidelity DNA Polymerase (2 U/µL) and 1 µL each of forward and reverse primers (10µM, but 5 µM in 2016). The forward primer, gITS7 (GTGARTCATCGA*R*TCTTTG) differed with one nucleotide code (*R* → *A*) from the one used for year 2 and fresh wood samples (fITS7) (Ihrmark et al., 2012). Reverse primer for both years was ITS4 (TCCTCCGCTTATTGATATGC (White, Bruns, Lee, & Taylor, 1990)). PCR was run on an GeneAmp® PCR System 2700 (Life Technologies, Applied Biosystems) with following settings: initial denaturation at 98°C for 30 s, then 30 cycles of 98°C/10 s denaturation, 56°C/30 s annealing and 72°C/15 s elongation, ending with 10 min elongation at 72°C. Amplifications were cleaned with Wizard® SV Gel and PCR Clean-Up System (Promega, Madison, WI, USA) as recommended by the manufacturers but with a longer centrifuging step of Membrane Wash Solution to remove excess ethanol.

A qualitative assessment was made of the cleaned PCR products with electrophoresis on a 1% agarose gel, ordering each sample in one of three categories based on band strength. The average amount of DNA for each category was estimated from a random selection and quantified with Qubit™ dsDNA HS Assay Kit on a Qubit® 2.0 Fluorometer (Life Technologies, Invitrogen). All samples were then normalized according to these categories before pooling them in three libraries. The libraries were purified with Agencourt AMPure XP magnetic beads (Beckman Coulter, CA, USA). Quality control was also done with Qubit. The libraries were barcoded with Illumina adapters, indexed with 30% PhiX and sequenced in an Illumina Miseq (Illumina, San Diego, CA, USA) lane with 2 × 300 bp paired-end reads at StarSEQ (StarSEQ GmbH, Mainz, Germany).

### Bioinformatic analyses

The sequencing runs resulted in two datasets: I. Two libraries from year 2 and fresh wood samples that consisted of 60 822 032 sequence reads. II. Three libraries from year 5 samples that consisted of 28 249 006 reads. The two datasets were demultiplexed in separate steps using CUTADAPT v2.7 (Martin, 2011). Tags, forward and reverse primers were cut simultaneously and reads less than 100 bp were discarded (dataset I: 6 294 571 reads, dataset II: 3 293 456 reads). Ten samples with low read numbers were removed (I: 2 samples, II: 8 samples). The reads were denoised, dereplicated, checked for chimeras and merged with DADA2 v1.12 (Callahan et al., 2016) in R v3.5.1 (Team, 2021). Before denoising, low quality samples were filtered with ‘filterAndTrim’ (Supp. Material 2 for details), but with optimised parameters for dataset I to retain more sequences without changing the read length characteristics (Figure S2). The reads were merged while allowing overhangs with ‘mergePairs’. Putative chimeric sequences were checked and removed with ‘removeBimeraDenovo’, specifying a more relaxed abundance distance threshold from parental sequences for dataset II. After combining the two datasets, the resulting table contained 26 395 685 reads and 7 626 amplicon sequence variants (ASVs) from 560 samples. We used the ITSx algorithm (v1.1b1; Bengtsson-Palme et al., 2013) to remove conserved regions adjacent to ITS2 and non-fungal ASVs. Then, with VSEARCH v2.9.1 (Rognes, Flouri, Nichols, Quince, & Mahé, 2016), the reads were dereplicated and clustered with 97% similarity using a distance-based greedy clustering, resulting in 1 668 operational taxonomic units (OTUs). These were then curated with the LULU algorithm (Frøslev et al., 2017) to avoid potential oversplitting, ending up with 1 287 OTUs. We assigned taxonomy to 1 126 OTUs (88.13%) with a BLAST+ v 2.8.1 search (evalue 1 × 10^−4^, max_target_seqs 1) against the UNITE+INSD v8.2 database of fungal sequences (Abarenkov et al., 2020). Information on ecological guilds were annotated to 758 OTUs (58.81%) using funGUILD (Nguyen et al., 2016) as of February 2020.

### Statistical analyses

All analyses and figure preparation were carried out in R v4.1.1 (Team, 2021). Rarefaction and richness estimations were performed with package PHYLOSEQ (McMurdie & Holmes, 2013) and ordinations with package VEGAN v2.5-7 (Oksanen et al., 2013).

Invertebrate exclusion, landscape and log section were used as categorical, while and bark resource quality and initial fungal community were used as continuous, explanatory variables to explain successional fungal community composition, OTU richness and wood density (Table S3). Bark variables were used as average values per tree (n = 17), while wood variables stemmed from the closest wood disc that had been sampled between logs at each tree (Figure S1). All continuous variables were transformed to zero skewness (Økland, Økland, & Rydgren, 2001) to approximate normality of errors, i.e. log-transformation of right-skewed and exponential transformation of left-skewed variables, then standardized to a 0-1 scale (Table S3). The initial fungal community was identified by DNA analysis and described by ordination – specifically, the two first axes of a detrended correspondence analysis (DCA1 and DCA2) (see Supp. Material 3.1). Wood and bark secondary metabolites from fresh wood were used as the total sum of all phenolic groups (phenolic acids, salicylates and flavonoids), as identified by T Birkemoe et al. (2021). Additional variables of resource quality were: bark MeOH-soluble and -insoluble condensed tannins; and wood carbon and nitrogen.

Negative and technical replicates were checked and are described in Supp. Material 3.2. For each pair of technical replicates, we retained the sample with most reads. All analyses were performed on a filtered (OTUs >10 reads) and rarefied dataset of samples from year 2 and 5 (Supp. Material 3.3). We removed 26 samples from year 2 and 11 samples from year 5 because they were either below the subsampling depth (14 407 reads) or not present in both years. This resulted in a dataset of 773 fungal OTUs from 426 samples originating from 213 aspen logs.

Fungal community composition of year 2 and 5 was explored with four-dimensional global nonmetric multidimensional scaling (gNMDS) (Kruskal, 1964a, 1964b) on a Bray-Curtis dissimilarity matrix. Ordination settings and evaluation are described in Supp. Material 4.1. The samples were balanced between the two years to consider displacement of the logs in ordination space. The axes were sorted by most variation explained and scaled in half-change units of compositional turnover (Faith, Minchin, & Belbin, 1987). We assessed which variables were driving the variation in fungal community composition of year 2 and 5 in three different ways: (1) linear mixed models (LMMs) of gNMDS axes, (2) hypothesis testing with partial constrained ordination and, (3) fitting variables onto the gNMDS with ‘envfit’. (1) LMMs were fitted to each of the four main axes identified by gNMDS using package LME4 (Bates, Mächler, Bolker, & Walker, 2014) with site as random effect. Models were chosen after forward selection with second-order AIC (AICc, Burnham and Anderson (2002); package MuMIn (Barton, 2020)). Degrees of freedom and p values were calculated with Satterwaithe’s approximation from the LMERTEST package (Kuznetsova, Brockhoff, & Christensen, 2017) and α set to 0.05. Additionally, we used Tukey’s HSD test to see if the invertebrate exclusion treatment levels were different along the gNMDS axes (Tukey, 1949). (2) Partial constrained ordination was performed with canonical correspondence analysis (CCA; Ter Braak (1986)), then forward selection of explanatory variables and a Monte Carlo permutation test (perm. = 999; Legendre, Oksanen, and ter Braak (2011)). The *p* value was used as selection criterion, or or F value if *p* values were equal between competing models. Model selection was done on the dataset from year 2, year 5 and both datasets combined. In the final models, we used inertia units to compare the variation explained by each variable. As the final models contained at least seven variables and the shared variation between variables was negligible, we chose to present variation explained by each variable independent of all others. (3) Variables that were congruent with both tests were fitted to the gNMDS with the ‘envfit’ function from VEGAN. This is based on factor averages and vectors (for continuous variables) and the significance of the variables fit was tested (perm. = 999).

To test whether wood density (g/cm^3^) or fungal OTU richness in year 2 and 5 were related to initial resource quality, assembly history, log section or landscape, we applied LMM as described above. Model assumptions were deemed satisfactory except a very severe violation of linearity of fixed effect residuals (Figure S5.1). This was because the bark and fresh wood values were derived from fewer samples than the wood density and fungal OTU samples. To provide more certainty to these results, we therefore also tested OTU richness and wood density of year 2 and 5 in linear regressions with aggregated datasets (i.e. n = 17 for bark samples and n = 50 for wood samples; Supp. Material 5).

## Results

### Taxonomic and functional groups of OTUs

There were 773 OTUs, of which 537 occurred in year 2 and 422 in year 5. Most OTUs were Ascomycota (61.1%) and 27.6% were Basidiomycota. Helotiales was the most common order (116 OTUs), followed by Agaricales (36) and Polyporales (34). Around half of the reads from year 2 (52%) and a third (32%) from year 5 were from OTUs annotated as «Wood saprotrophs». The functional distribution of OTUs across both years were: 411 saprotrophs (of which 118 were wood saprotrophs), 156 symbiotrophs (e.g. mycorrhiza), 153 pathotrophs and 243 that were not assigned to guilds. Note that many OTUs are assigned to more than one guild, e.g. pathotroph-saprotroph. The initial fungal community from fresh wood consisted of 375 OTUs of which 68.8% were Ascomycota and 18.9% Basidiomycota. The functional distribution was: 171 saprotrophs (49 wood saprotrophs), 87 symbiotrophs, 83 pathotrophs and 129 that were not assigned to guilds. Note that while it is possible that this may include airborne fungal spores that arrived during sampling, it still reflects the initial fungal community.

### Fungal community composition

We identified four compositional gradients of the fungal community from year 2 to 5 that were interpretable in terms of temporal variation, spatial variation, initial resource quality and assembly history. Specifically, the main compositional gradient (gNMDS1) separated the fungal communities from year 2 to year 5, while also being related to spatial variation and wood phenolic acids (Figure 1, Table S4.2). gNMDS3 was related to the initial fungal community (DCA1) and wood carbon (Figure S4.4, Table S4.2).The second and fourth axes formed gradients related to the initial fungal community (DCA2) and resource quality: from higher amounts of wood carbon to bark flavonoids in gNMDS2 (Figure 1), and from higher amounts of wood nitrogen to bark MeOH-soluble condensed tannins in gNMDS4 (Figure S4.4, Table S4.2). The second axis showed an invertebrate-induced priority effect, with a gradient from caged to positive control logs (Figure 1). Although the variable was not included in the LMM explaining variation in gNMDS2, a Tukey HSD test showed significant dispersion in plot scores between caged and positive control logs along this axis (*p* = 0.047; Figure S4.5).

**Figure 1.**
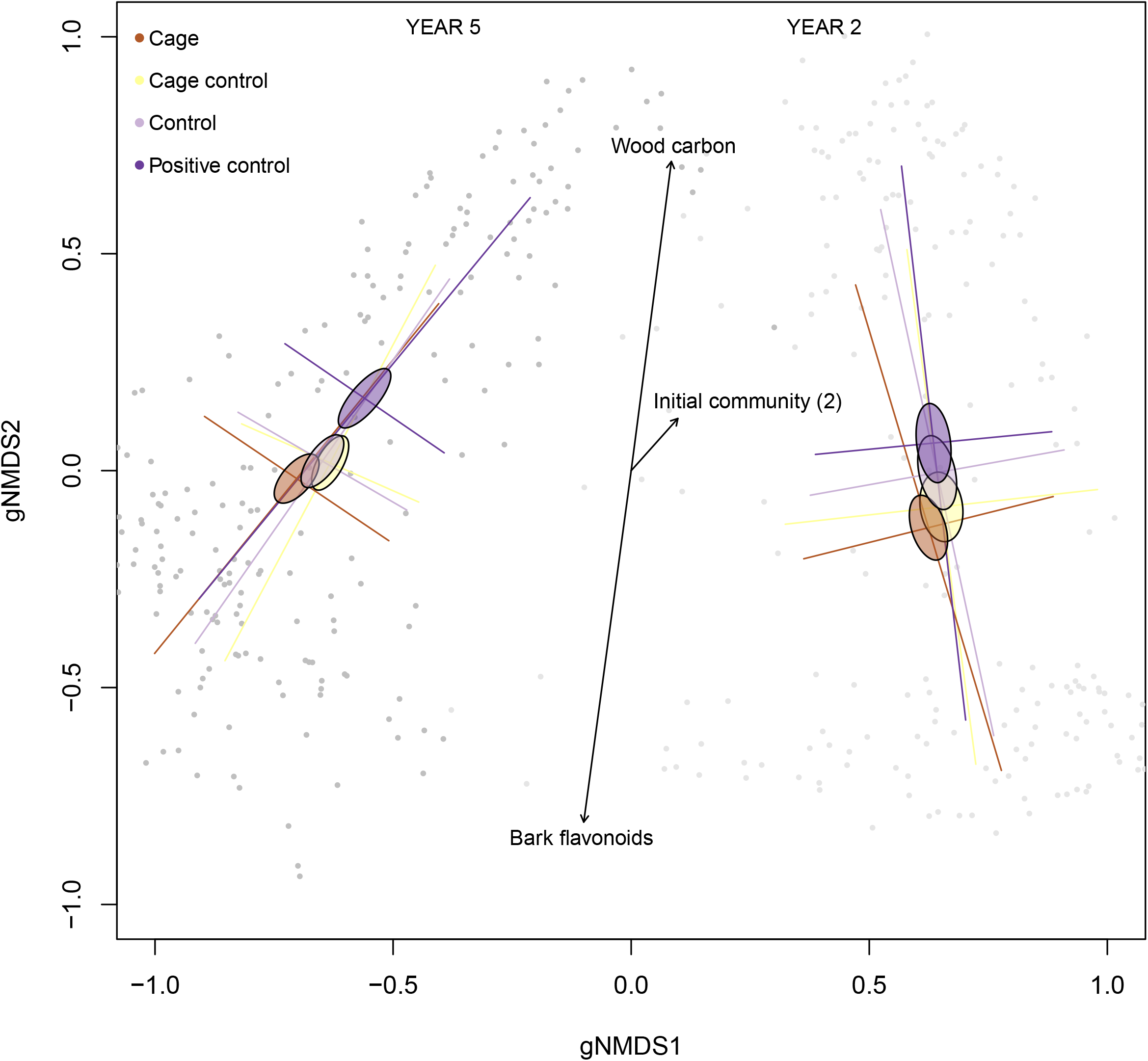
Ordination biplot of the first and second compositional gradients of fungal OTUs from aspen logs (n = 424) in year 2 and 5 after tree felling (gNMDS of Bray-Curtis dissimilarities; stress = 0.14). Coloured ellipses display the standard error of treatment levels, while error bars display the standard deviation. Vectors are fitted with the ‘envfit’ function in package VEGAN and are based on a forward selection of linear mixed models of each gNMDS axis. Light grey dots represent plot scores of year 2 and dark grey of year 5.

From the model selection with constrained ordination of both years, temporal variation accounted for 37.3% of the explained variation, followed by invertebrate exclusion (11.4%) and variation between landscapes (8.7%) (Table S4.3). Invertebrate exclusion was the most important initial condition explaining the fungal community composition in year 2 and year 5 (Table 1, 2). The relative importance of invertebrate exclusion increased from year 2 (10% of explained variation; Figure S4.6a) to year 5 (25%; Figure S4.6b). Combined, the secondary metabolites contributed to around half of the explained variation in year 2 and 5 (Figure S4.6a, b). All initial conditions were deemed important in explaining fungal community composition in year 2, while invertebrate exclusion, flavonoids, phenolic acids and nitrogen were important in year 5 (Table 1, 2).

**Table 1.** Variation partitioning with constrained ordination (CCA) of fungal community composition in aspen logs in year 2 after tree felling (n=213). Variables are based on a forward model selection with *p* as selection criterion. Inertia of constrained variables are modelled alone without conditioning variables to show variation explained independent of other variables.

| Variable | Inertia | Prop. explained |
| --- | --- | --- |
| Landscape | 0.1335 | 0.0200 |
| Invertebrate exclusion | 0.1142 | 0.0171 |
| Bark flavonoids | 0.0879 | 0.0131 |
| Wood phenolic acids | 0.0849 | 0.0127 |
| Bark sol. tannins | 0.0805 | 0.0120 |
| Wood salicylates | 0.0788 | 0.0118 |
| Wood flavonoids | 0.0750 | 0.0112 |
| Wood carbon | 0.0702 | 0.0105 |
| Initial fungal community (1) | 0.0679 | 0.0101 |
| Bark phenolic acids | 0.0651 | 0.0097 |
| Initial fungal community (2) | 0.0641 | 0.0096 |
| Bark insol. tannins | 0.0640 | 0.0096 |
| Log section | 0.0586 | 0.0088 |
| Wood nitrogen | 0.0504 | 0.0075 |
| Unconstrained | 5.7361 | 0.8574 |
| Total | 6.6900 | - * |

\* Inertia of constrained and unconstrained axes does not add up to the total inertia because 2.11% of the variation is shared between one or several variables.

**Table 2.** Variation partitioning with constrained ordination (CCA) of fungal community composition in aspen logs in year 5 after tree felling (n=213). Variables are based on a forward model selection with *p* as selection criterion. Inertia of constrained variables are modelled alone without conditioning variables to show variation explained independent of other variables.

| Variable | Inertia | Prop. explained |
| --- | --- | --- |
| Invertebrate exclusion | 0.3071 | 0.0173 |
| Landscape | 0.2049 | 0.0116 |
| Bark phenolic acids | 0.1681 | 0.0095 |
| Bark flavonoids | 0.1395 | 0.0079 |
| Wood phenolic acids | 0.1314 | 0.0074 |
| Wood flavonoids | 0.1272 | 0.0072 |
| Wood nitrogen | 0.1027 | 0.0058 |
| Unconstrained | 16.5813 | 0.9352 |
| Total | 17.7301 | - * |

\* Inertia of constrained and unconstrained axes does not add up to the total inertia because 0.18% of the variation is shared between one or several variables.

### Wood density

Bark phenolic acids and MeOH-insoluble condensed tannins, as the only included variables after model selection, had a positive effect on wood density (Table 3). The relative contribution of bark tannins was stronger in year 5 than 2. Phenolic acids only explained wood density in year 2, but this effect was not consistent in models that took into account linearity of fixed effect residuals (Supp. Material 5).

**Table 3.**
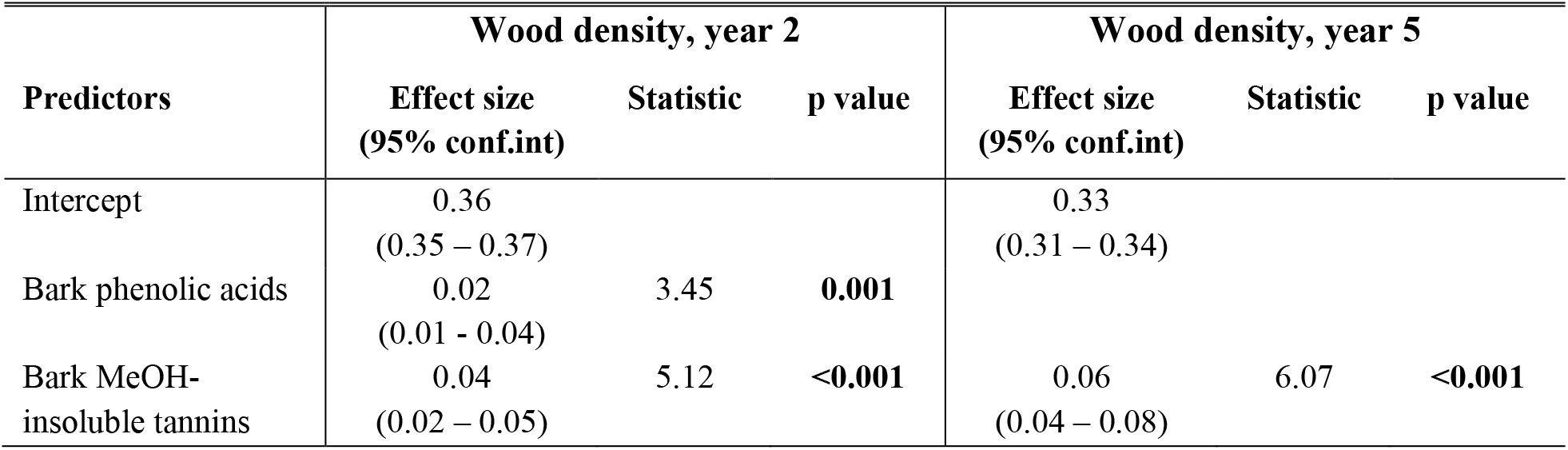
Linear mixed models of initial conditions explaining variation in aspen wood density in year 2 and 5 after tree felling. The models are based on a forward model selection (AICc) with Site as random effect.

| Predictors | Wood density, year 2 |  |  | Wood density, year 5 |  |  |
| --- | --- | --- | --- | --- | --- | --- |
|  | Effect size<br>(95% conf.int) | Statistic | p value | Effect size<br>(95% conf.int) | Statistic | p value |
| Intercept | 0.36<br>(0.35 – 0.37) |  |  | 0.33<br>(0.31 – 0.34) |  |  |
| Bark phenolic acids | 0.02<br>(0.01 - 0.04) | 3.45 | **0.001** |  |  |  |
| Bark MeOH-insoluble tannins | 0.04<br>(0.02 – 0.05) | 5.12 | **<0.001** | 0.06<br>(0.04 – 0.08) | 6.07 | **<0.001** |

### Fungal OTU richness

Wood carbon was negatively correlated with fungal OTU richness in year 2 (Figure 2, Table S6). Also, richness was estimated to be higher in mid, rather than end, log sections (p = 0.019) and in Østmarka, rather than Nordmarka landscape (p = 0.039). Wood phenolic acids positively affected fungal OTU richness in year 5 (p = 0.013). Bark flavonoids positively affected fungal OTU richness both years, but the effect was not significant (p = 0.07).

**Figure 2.**
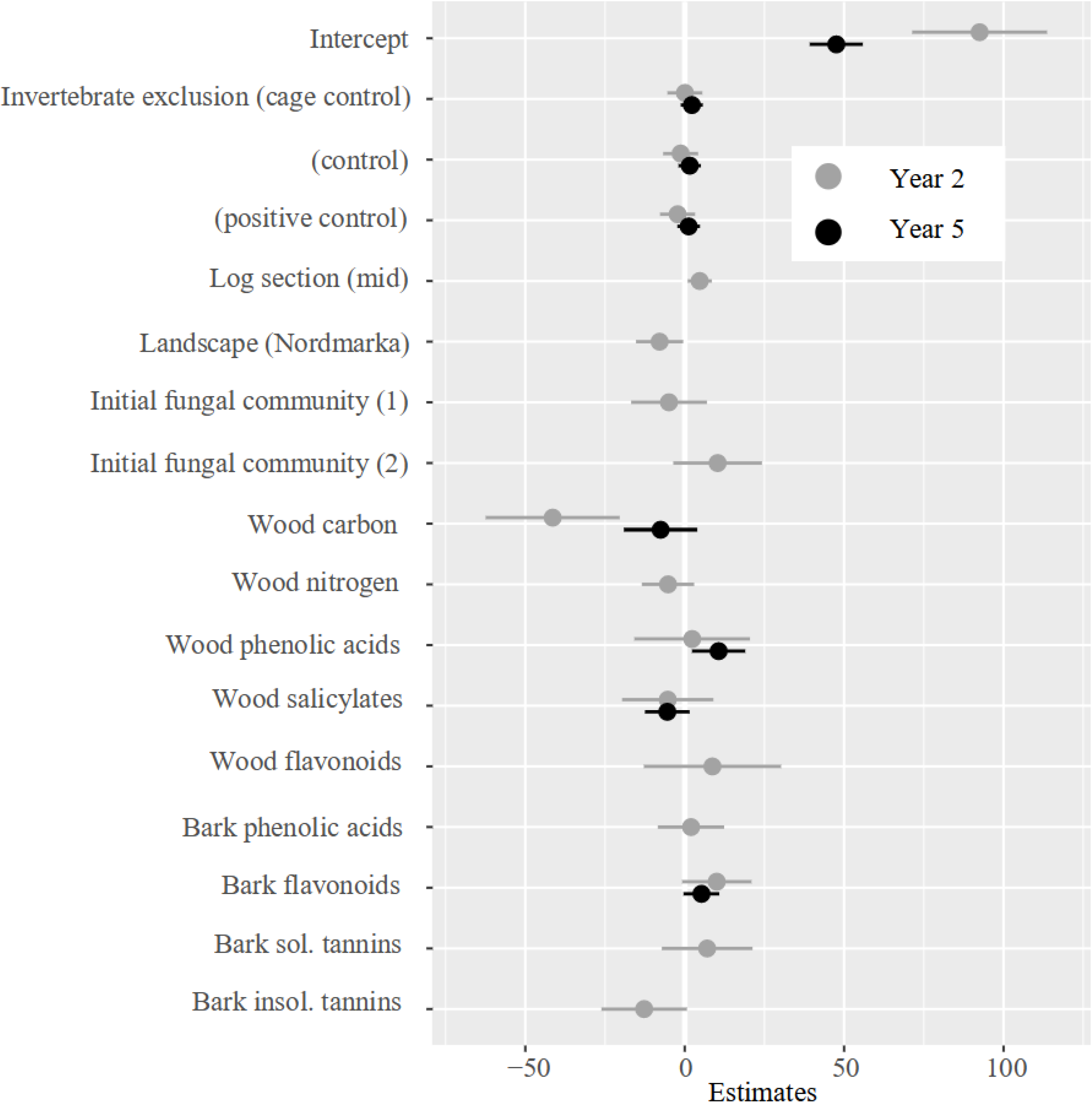
Linear mixed model effect sizes of initial conditions explaining fungal OTU richness in aspen logs in year 2 and 5 after tree felling. Based on a forward model selection with Site as random effect. Effect sizes that does not overlap with zero are significant (α = 0.05). Reference levels in intercept are: Invertebrate exclusion (cage), Log section (end) and Landscape (Østmarka).

## Discussion

By combining DNA metabarcoding with chemical analyses in an experimental field study of dead wood of aspen, we have shown that exclusion of invertebrates had contingent effects on the fungal community for at least 3 years after exclusion ceased. However, the initial fungal community present in fresh wood was only important after 1.5 years of succession. Moreover, intraspecific variation in initial bark and wood secondary metabolites, carbon and nitrogen affected not only fungal community composition, but also wood decomposition rates.

### Secondary metabolites, carbon and nitrogen structure fungal communities

After 1.5 and 4.5 years of succession, we identified compositional gradients of dead wood fungi that were related to both the initial fungal community and resource quality. The gradient spanned from higher amounts of carbon or nitrogen in one end, to higher amounts of secondary metabolites in the other end. Specifically, concentration of flavonoids and phenolic acids from bark and wood affected fungal community composition and diversity.

Fungi in dead wood have evolved different life history strategies based on the amount of stress, disturbance and competition they face within their environment (Lynne Boddy, 2001). Phenolic compounds, including flavonoids, from wood can cause intracellular stress by directly disrupting the degradative capacities of dead wood fungi and even destroying the fungal cell (Valette et al., 2017). Indeed, phenolics isolated from bark and wood of American aspen (*Populus tremuloides*) inhibited growth and spore germination of a tree canker-causing fungus, *Entoleuca mammata* (Flores & Hubbes, 1979, 1980; Hubbes, 1962, 1969). The gradients we identify that structure fungal communities in relation to resource quality may reflect strategies to tolerate stress and competition for territory, i.e. combat. For instance, in resources with higher concentrations of secondary metabolites, a combative species might have lower net energy intakes or use more energy on maintenance, thereby reducing its combative abilities relative to that of a stress-tolerant species (Crowther et al., 2014; Schoener, 1973). Because of the harsh environment that latent fungal communities face when the tree is alive, many of these species are stress-tolerant (Lynne Boddy & Heilmann-Clausen, 2008; Rayner & Boddy, 1988). Indeed, the trade-off between combative ability and stress tolerance in dead wood fungi has been strongly asserted (Lynne Boddy & Heilmann-Clausen, 2008; Crowther et al., 2014; Maynard et al., 2019; Treseder & Lennon, 2015). Environmental gradients in resource quality can increase the number of ecological niches that are available in the wood and allow more fungal species to coexist. This is suggested by the positive effect that wood phenolic acids and bark flavonoids had on fungal OTU richness in our study. Even though we have not measured the combative ability of fungal species, nor classified the fungal community according to life history strategies, our results show that gradients of resource quality structure successional fungal communities in dead wood.

### Initial fungal community has transient effects on fungal community development

The initial fungal community was an important driver of community composition after 1.5 years, but not after 4.5 years, of succession. Song, Kennedy, Liew, and Schilling (2017) found an effect of the initial fungal community on later fungal colonizers in isolated mesocosms. However, their study ended after only 6 months of succession and they excluded natural colonizers. Field studies agree with our findings, that latent fungal communities have similar short-term priority effects in early stages of succession, but that these effects attenuate over time (Chapela & Boddy, 1988a; Cline et al., 2018; van der Wal, Klein Gunnewiek, Cornelissen, Crowther, & de Boer, 2016). This can be because the initial fungal colonizers are replaced over time by secondary colonizers with highly combative abilities (Chapela, Boddy, & Rayner, 1988; van der Wal et al., 2016) and highlights the importance of long-term observations in studies on dead wood communities. Secondary colonizers arrive in the dead wood via air-borne or animal-vectored spores, or from the soil as cords or rhizomorphs (Lynne Boddy & Heilmann-Clausen, 2008). As we placed all logs on plastic sheets to exclude soil invertebrates from the logs during the first two summers, this would have also excluded fungi that colonize from the soil with mycelial cords. Cord-forming fungi are strong competitors (Hiscox et al., 2018; Hiscox, Savoury, Vaughan, Müller, & Boddy, 2015; Rayner & Boddy, 1988) and this could explain why the effect we found of the initial fungal community was only transient; when the plastic sheets were removed after 1.5 years, secondary colonizers, such as cord-forming fungi, may have replaced the initial community via competitive exclusion.

### Invertebrate-induced priority effects after 4.5 years of fungal community development

We detected a strong invertebrate-induced priority effect on fungal community composition, even after 4.5 years of succession, although the experimental set-up excluding invertebrates was removed after 1.5 years. Many invertebrates help fungi to establish in dead wood substrates, for example bark beetles that vector ascomycete mutualists (Klepzig & Six, 2004) and other rot fungi (Harrington et al., 1981; Persson, Ihrmark, & Stenlid, 2011). Dispersal of fungi by non-mutualistic beetles to dead wood also seems to be common, although the importance for fungal colonization and population dynamics remains unknown (Rannveig M Jacobsen, Kauserud, Sverdrup-Thygeson, Bjorbækmo, & Birkemoe, 2017; Seibold et al., 2019). On the other hand, invertebrates can reduce fungal growth (A’Bear, Jones, et al., 2014) or alter the outcomes of competitive interactions between fungi (Crowther, Boddy, et al., 2011; Rotheray, Chancellor, Jones, & Boddy, 2011), as shown in soil microcosms by mycelial grazers. However, as indicated by studies on the woodlouse (Isopoda) *Oniscus asellus*, the effects of fungivory may be less in the field than in laboratory experiments (A’Bear, Boddy, Kandeler, Ruess, & Jones, 2014; Crowther & A’Bear, 2012; Crowther, Boddy, et al., 2011). This makes our results even more intriguing. Although we cannot know the underlying causes behind the invertebrate-induced priority effects demonstrated in our study, long-term studies show that beetles can have variable, but significant, effects on later fungal colonizers (Rannveig Margrete Jacobsen et al., 2015; Weslien et al., 2011). By providing evidence of invertebrate-induced priority effects at the community scale, we add crucial knowledge to the importance of invertebrates in affecting fungal community development through altered assembly history.

### Bark secondary metabolites impede wood decomposition

High concentration of bark phenolic acids and MeOH-soluble condensed tannins impaired wood decomposition rates, but only tannins had an effect 4.5 years after tree felling. Simple molecular compounds, like phenolic acids, are targeted during early stages of litter decomposition (Chomel et al., 2016; Moorhead & Sinsabaugh, 2006). Tannins, on the other hand, are polymers of repeating phenolic units and are therefore more difficult to break down. Further, condensed tannins complex with proteins and even with fungal chitin (Adamczyk, Sietiö, Biasi, & Heinonsalo, 2019), thus altering the enzymatic activity of microbes and may as such impede decomposition considerably (Adamczyk, Adamczyk, Smolander, Kitunen, & Simon, 2018; Kraus, Dahlgren, & Zasoski, 2003; Smolander, Kanerva, Adamczyk, & Kitunen, 2012). In a study on four tropical tree species, Loranger et al. (2002) found that simple phenolic compounds controlled early stages of leaf litter decomposition, while later stages were correlated by tannin concentrations. Although we only have initial measurements of secondary metabolites, we suspect that the concentrations of phenolic acids in aspen logs decreased more than tannins during the study period, explaining the lingering effect of tannins after 4.5 years.

In a study using the same samples as here, Rannveig M Jacobsen et al. (2018) found, in contrast to us, a significant difference between logs where invertebrates had been excluded and control logs, after 1.5 years of succession. The essential difference between our two models was that we did not fit the tree identity nor the log’s position on the trunk as random effects, because we aimed to use initial secondary metabolite, carbon and nitrogen content to explain this variation. The diverging results between our studies indicate that there was considerable variation between or within trees that we did not manage to account for. Indeed, many other tree qualities may govern wood decomposition, such as bark anatomy (Dossa et al., 2018), wood water content (Rayner & Boddy, 1988; Venugopal et al., 2017), or the identity of fungal (Rayner & Boddy, 1988; Venugopal et al., 2017), invertebrate (Seibold et al., 2021; Ulyshen, 2016), or even bacterial (Johnston, Boddy, & Weightman, 2016), communities.

The altered effect of several variables from 1.5 to 4.5 years in our study underlines the importance of long-term observations in dead wood systems. Many earlier studies that have linked priority effects to wood mass loss only lasted for 1 year (Dickie et al., 2012; Leopold et al., 2017; Song et al., 2017), although wood decomposition might take decades (Stokland et al., 2012). It appears that the link between community development and functioning is far from obvious. Over 4.5 years, we found that resource quality, but not assembly history, were drivers of wood decomposition rates. However, altered assembly history through invertebrate exclusion controlled fungal community development, even 3 years after exclusion had ceased.

## Acknowledgements

We would like to thank Tone Aasbø Granerud and David Arnott for help with sawdust sampling. Thanks to Mina-Johanne Tangnæs, Martine Andelic, Malin Stapnes Dahl and Sundy Maurice for help in the lab. A special thanks to Luis Morgado for help in the lab and for doing bioinformatics. Thank you, Magnus Nygård Osnes and Rune Halvorsen, for taking the time to discuss statistics and ordination.

## Data accessibility

The raw sequence reads are deposited in NCBI SRA SUB10694654. The OTU table, metadata, mapfiles and R script for reproducing figures and analyses in the manuscript are deposited in Dryad (doi:10.5061/dryad.zcrjdfndj).

## Author contributions

LFL performed laboratory work, bioinformatic and statistical analyses, and led the writing of the manuscript; TB conceived, designed and coordinated the project and performed sampling; RMJ conceived and designed the project, performed sampling, laboratory work and contributed to statistical analyses; AST and HK designed the project; LN contributed to chemical analyses. All authors contributed to interpretation and writing of the manuscript, and gave final approval upon publication.

