## Supplementary Material for "Invertebrate-induced priority effects and secondary metabolites control fungal community composition in dead wood"


1. **Study design** **p. 2**
2. **Bioinformatics analyses** **p. 3**
3. **Data preparation** **p. 4**
   - **3.1. Initial fungal community** **p. 5**
   - **3.2. Negative and technical replicates** **p. 7**
   - **3.3. Rarefaction and read depth** **p. 7**
4. **Multivariate analyses** **p. 9**
   - **4.1. gNMDS ordination** **p. 9**
   - **4.2. Partial constrained ordination** **p. 13**
   - **4.3. 'Envfit'** **p. 14**
5. **Model diagnostics** **p. 17**
   - **5.1 Fungal OTU richness** **p. 17**
   - **5.2 Wood density** **p. 19**
6. **Model of fungal OTU richness** **p. 22**

**References** **p. 23**

#### 1. Study design

As there were more samples from the experimental logs than from fresh wood (Figure S1), we linked (i.e. duplicated) the values from fresh wood discs (3, 7 and 11 in Figure S1) to the nearest experimental logs. This meant that fresh wood variables from wood disc 3 of, for example tree A, would be the value of log 1, 2, 4 and 5 of tree A (Figure S1, in red). Values from wood disc 7 linked to log 6, 8 and 9 (Figure S1, in black); values from wood disc 11 linked to log 10, 12 and 13 (Figure S1, in blue), and; values from wood disc 15 (not all trees) linked to log 14, 16 and 17. There were only two wood discs from tree Q: disc 6 duplicated values for logs from 1-9 and disc 14 for logs 10-13.

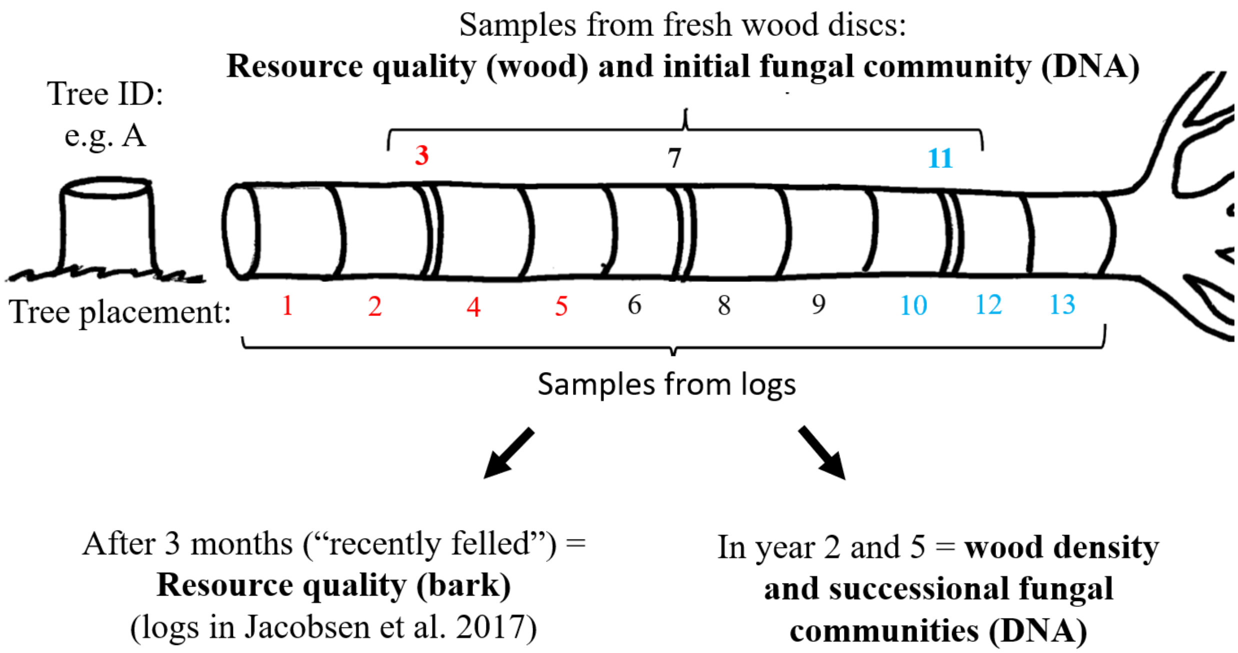


**Figure S1.** Sample design for fungal OTUs (initial and successional communities) and resource quality of wood and bark. All samples were from the same 17 aspen trees, but they differed as to: when they were sampled, which tree (tree ID) and which position on the tree (tree placement) the log came from. Samples from fresh wood discs (wood resource quality and initial fungal community) were linked to the nearest wood log to fit them in models explaining variation in successional fungal OTU community composition, richness and wood density. The linked discs and respective logs are given the same colours (red, black and blue).

#### 2. Bioinformatic analyses

The sequencing runs resulted in two datasets: I. Two libraries from year 2 and fresh wood samples that consisted of 60 822 032 sequence reads. II. Three libraries from year 5 samples that consisted of 28 249 006 reads. DADA2 ran separately for the two datasets because DNA isolation, amplification and sequencing had been done at different times. While most settings were the same between the two datasets, there was a need to optimise parameters during pre-denoising clean-up of year 2 samples to avoid losing too many reads (filterAndTrim, discarding reads with quality scores below 2, maximum expected errors 2 (dataset II) or 3 (dataset I), and maximum truncated read length 40 (dataset II) or 80 (dataset I)). The optimisation improved percentage of reads kept without disturbing the read length characteristics (Figure S2). We then dereplicated, error corrected, denoised (learnErrors: verbose = T, multithread = T, dada: err = 'learnErrors', pool = 'pseudo', multithread = T) and merged the datasets (mergePairs, allowing for overhangs). Putative chimeric sequences were checked and removed, specifying a more relaxed abundance distance threshold (i.e. 8) for parental sequences in dataset II (removeBimeraDenovo).

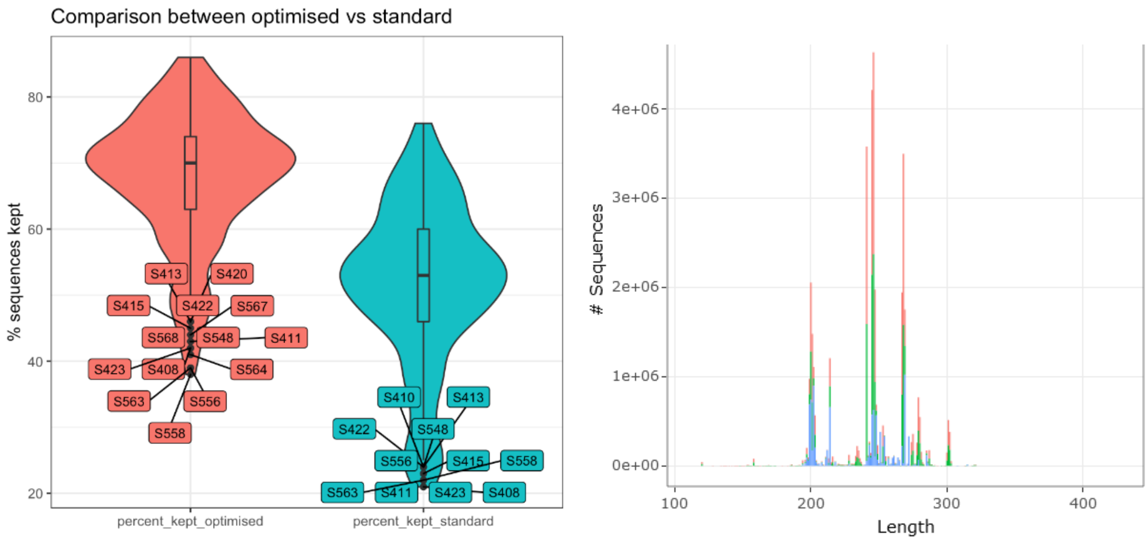


**Figure S2.** Comparison of performance with standard (blue) and optimized (red) settings during pre-denoising clean-up with 'filterAndTrim' for dataset I. With optimization, more reads were kept per sample (left) without changing the read length characteristics (right). Read lengths of dataset II in green.

### 3. Data preparation

**Table S3.** Overview of explanatory variables. Transformation tells how the continuous variables were transformed prior to analyses. The last column show the number of unique values, for continuous, or levels for categorical variables. Cont = continuous, of which some are groupings of phenolic compounds and represent the total sum of several individual compounds (described in Birkemoe et al. 2021).

| Variable | Type | Variable type | Transformation | n values/levels |
|---|---|---|---|---|
| Site | Random effect | Categorical | - | 30 |
| Landscape | Spatial variation | Categorical | - | 2, Østmarka/Nordmarka |
| Year | Temporal variation | Categorical | - | 2, year2/year5 |
| Log section | Spatial variation | Categorical | - | 2, end/mid |
| Invertebrate-induced priority effects | Initial condition | Categorical | - | 4, cage/cage control/control/positive control |
| Initial fungal community (1) | Initial condition | Cont; DCA1 | Logarithmic | 48 |
| Initial fungal community (2) | Initial condition | Cont; DCA2 | Exponential | 48 |
| Wood carbon | Initial condition | Cont; | Exponential | 53 |
| Wood nitrogen | Initial condition | Cont; | Logarithmic | 53 |
| Wood flavonoids | Initial condition | Cont; summed | Logarithmic | 53 |
| Wood phenolic acids | Initial condition | Cont; summed | Logarithmic | 53 |
| Wood salicylates | Initial condition | Cont; summed | Logarithmic | 53 |
| Bark flavonoids | Initial condition | Cont; summed | Logarithmic | 17 |
| Bark phenolic acids | Initial condition | Cont; summed | Logarithmic | 17 |
| Bark MeOH-soluble condensed tannins | Initial condition | Cont; | Logarithmic | 17 |
| Bark MeOH-insoluble condensed tannins | Initial condition | Cont; | Logarithmic | 17 |

#### 3.1 Initial fungal community

The samples from fresh wood that were analysed for DNA were included as explanatory variables of compositional gradients to represent the initial fungal community. First, the samples were filtered (removed OTUs with <10 reads) and rarefied at the minimum subsampling depth (1356 reads). The dataset was then subjected to a multiple parallel ordination procedure involving detrended correspondence analysis (DCA) (Hill 1979, Hill and Gauch 1980), three gNMDS ordinations (two-, three- and four-dimensional) and four different weighting functions – rarefied, log (x+1)-transformed rarefied, presence-absence data and clr-transformed (function 'transform' from package MICROBIOME; Lahti and Shetty (2018)). Weighting function and ordination methods were chosen following the criteria of van Son & Halvorsen (2014) and interpreting the ordinations with wood chemistry variables from the same samples. We ended up with DCA from rarefied data based on 48 samples (after removing five outliers) and 375 OTUs (Figure S3.1a). Only the first two axes of the DCA were interpreted as structure axes. Although the second axis showed a tongue effect, we still believed that it represented real-structure compositional data because:

1) the gradient was interpretable in terms of the log's position on the trunk (Figure S3.1b; 'tree_placement')
2) the most common OTU, annotated to *Brunnipila palearum*, was present on the flattened part of the tongue and species scores of other common OTUs were distributed along the first gradient (Figure S3.2)

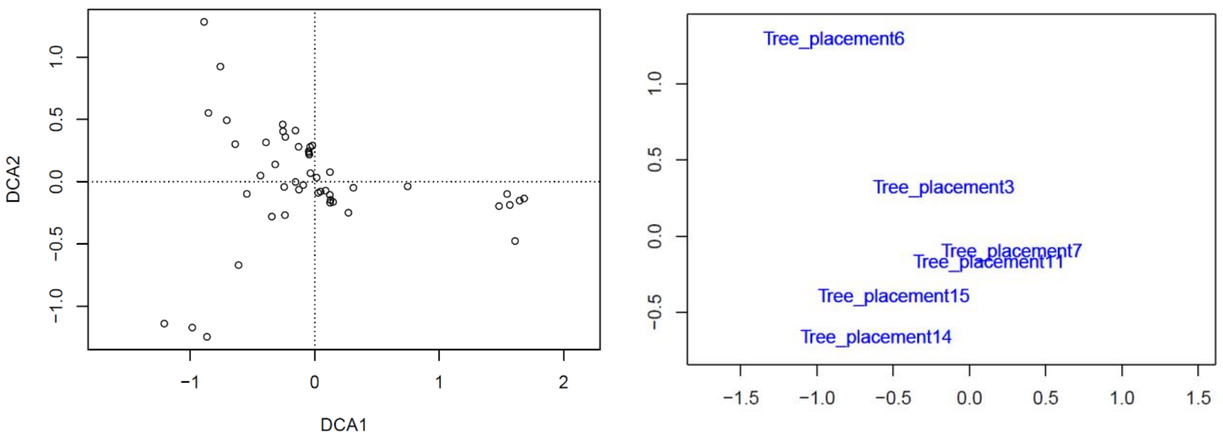


**Figure S3.1a-b.** Plot scores (a, left) of a detrended correspondence analysis (DCA) based on 375 OTUs from 48 samples of a initial fungal community in fresh aspen wood. (b, right) Factor averages of tree length levels fitted to DCA1 and 2 with the function 'envfit' ($p < 0.05$).

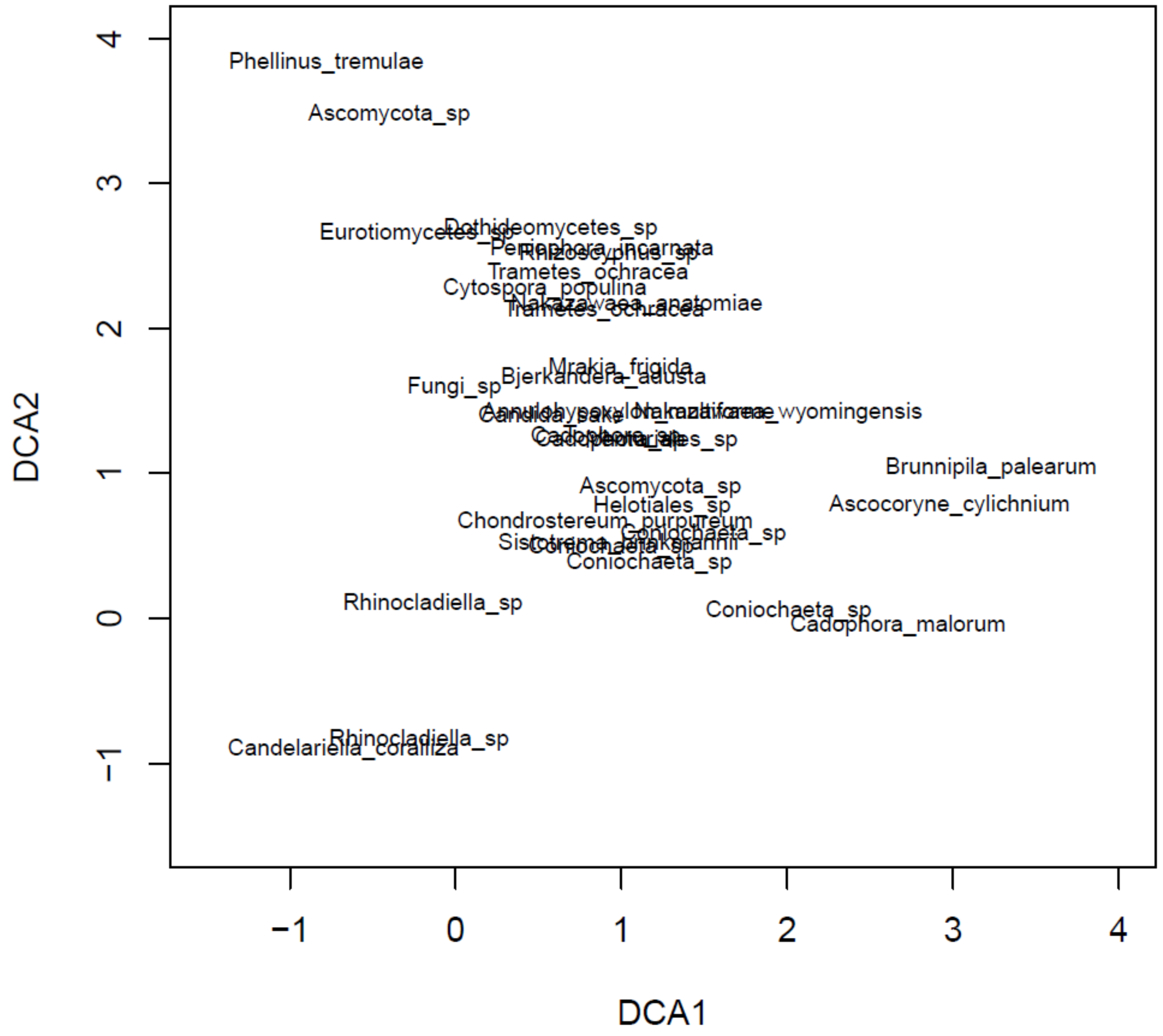


**Figure S3.2.** Species scores of a detrended correspondence analysis (DCA) based on 48 samples of a initial fungal community in fresh aspen wood. Only OTUs with more than 400 reads per sample are shown, i.e. the 33 most common initial fungi. Overlapping OTUs are: *Periophora incarnata* and *Rhizoschyphus* sp.; *Nakazawaea anatomiae* and *Trametes ochracea*; *Mrakia frigida* and *Bjerkandera adusta*; *Annulohypoxylon multiforme*, *Nakazawaea wyomingensis* and *Candida sake*; *Venturiales* sp., *Cadophora* sp. and *Cadophora* sp; *Coniochaeta* sp., *Sistotrema brinkmannii* and *Coniochaeta* sp.

##### 3.2 Negative and technical replicates

Six negative controls with high read numbers (3 from DNA isolation and 3 PCR negatives) and 60 technical replicates were examined for patterns of contamination. Exploration of negative replicates from year 5 showed substantial between-sample contamination, interpreted to have originated during PCR preparation or amplification (Figure S3.3). Subtraction of negative-sample reads from the other samples on a PCR-plate level resulted in even stronger differentiation between plates. Therefore, because technical replicate pairs clustered together, and because negative-sample reads were from real-data OTUs, we chose not to filter the dataset of negative-sample contamination.

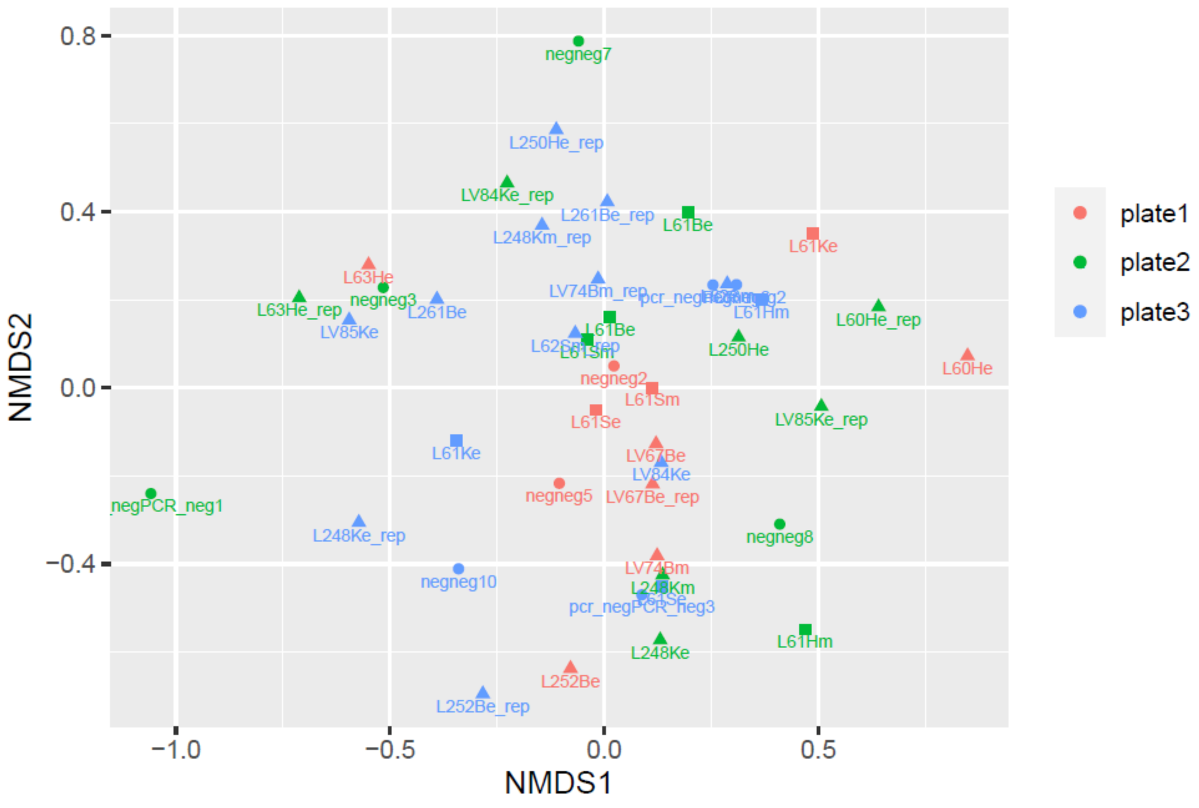


**Figure S3.3.** Ordination (gNMDS) of rarefied negative and technical replicates from fungal OTUs in aspen logs, year 5 after tree felling. Samples are coloured by PCR plate/library. Circles are negative replicates. Most replicates clustered well, but there were indications of divergence of samples based on plates.

##### 3.3 Rarefaction and read depth

All analyses were performed on a filtered (removed 356 OTUs with less than 10 reads) and rarefied dataset of samples from year 2 and year 5. We examined read depth between the two years (Figure S3.4) and rarefaction curves on a tree identity level (Figure S3.5). We set the rarefation subsampling depth to 14 407 reads per sample (i.e. minimum reads from year 2) to balance read depth between the two datasets.

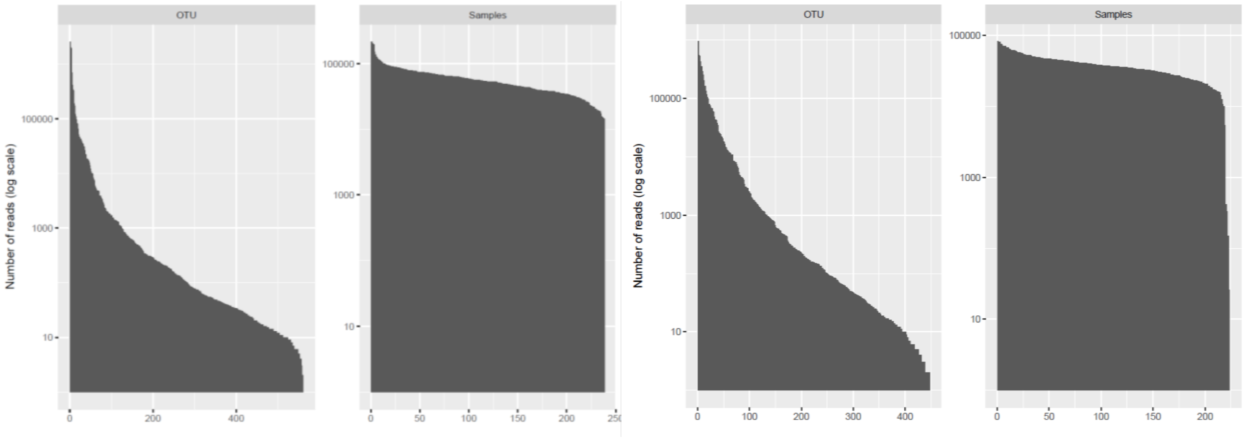


**Figure S3.4.** Read depth per OTU and per sample in year 2 (plots to the left) and year 5 (plots to the right). Y axis shows the number of reads on a logarithmic scale.

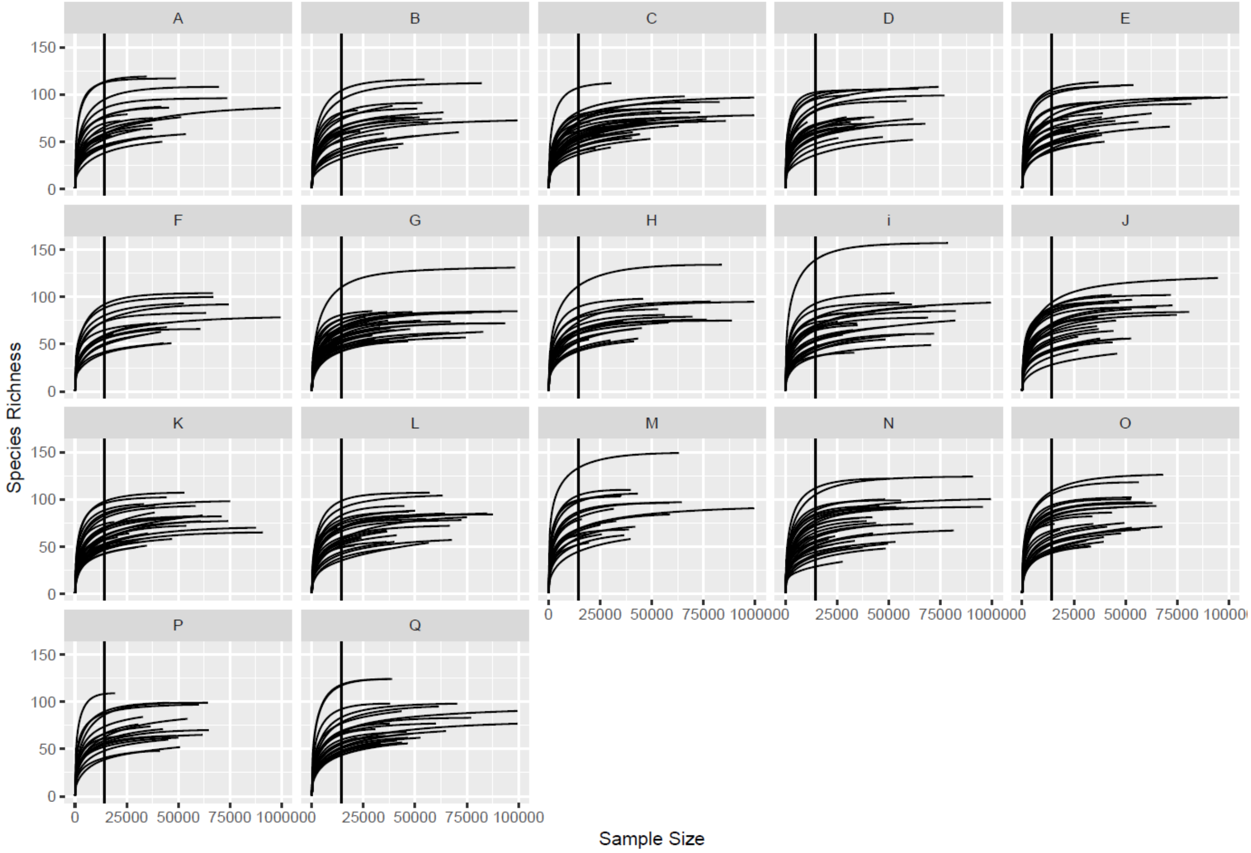


**Figure S3.5.** Rarefaction curves per tree individual. Vertical lines are drawn at the subsampling depth used for rarefaction (i.e. 14 407 reads).

### 4. Multivariate analyses

#### 4.1 gNMDS ordination

We ran a four-dimensional global non-metric multidimensional scaling (gNMDS) on fungal OTUs from year 2 and 5 after aspen felling with the following settings: Bray-Curtis dissimilarity measure, 100 initial starting configuration with maximum iteration at 200. Iterations would stop with stress or a gradient scale factor drop below 1 x $10^{-7}$, or stress change of 0.99999. We only accepted configurations if convergence had been reached.

We used Procrustes analysis to compare the 'best' configurations, i.e. with lowest stress values (Figure S4.1; test: perm = 999, $R^2 = 0.99$, p = 0.001) (Kent et al. 1979, Peres-Neto and Jackson 2001). We evaluated the 'best' configuration (stress = 0.14) with goodness-of-fit measure and Shepard plot. Then, we compared all the gNMDS axes with four axes of a DCA using Kendall's nonparametric correlation test (Kendall 1938) (Table S4.1) and principal component analysis (Figure S4.2; PCA; standardized and centered, correlation biplot scaling) (Pearson 1901; ter Braak and Prentice 1988)

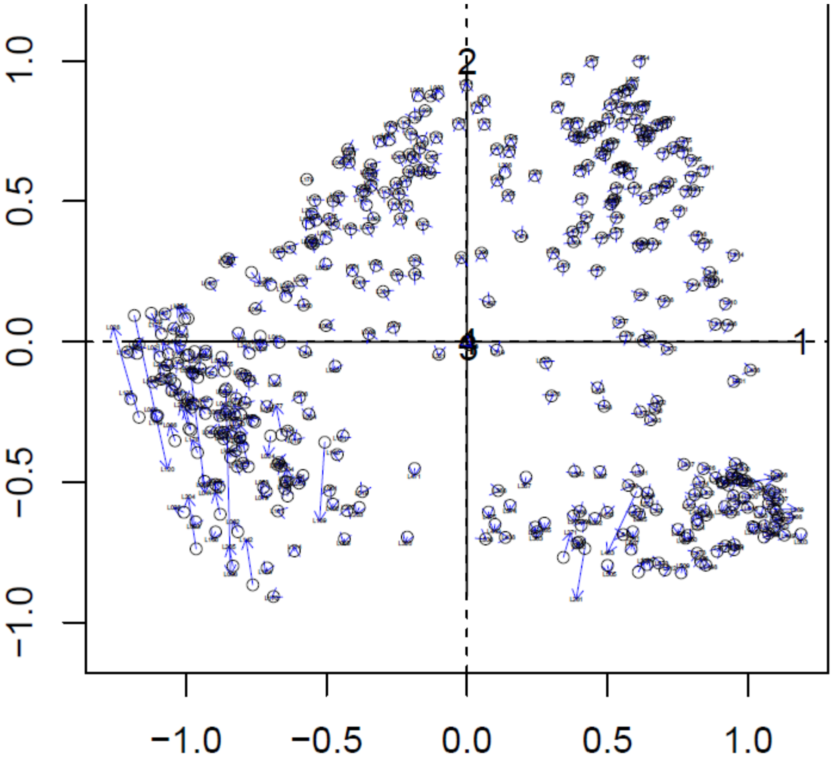


**Figure S4.1** Procrustes analysis evaluating the gNMDS configurations (showing axis 1 and 2, only) with lowest and second lowest stress values.

**Table S4.1**. Pair-wise comparisons of ordination axes (gNMDS and DCA) with Kendall's rank correlation test. All correlations were significant with the z statistic and 999 permutations.

| gNMDS axis | DCA axis | Kendall's τ |
|---|---|---|

| | | |
|---|---|---|
| gNMDS1 | DCA1 | **0.743** |
| gNMDS2 | DCA2 | **0.262** |
| gNMDS2 | DCA4 | **0.464** |
| gNMDS3 | DCA2 | **0.253** |
| gNMDS4 | DCA3 | **0.389** |

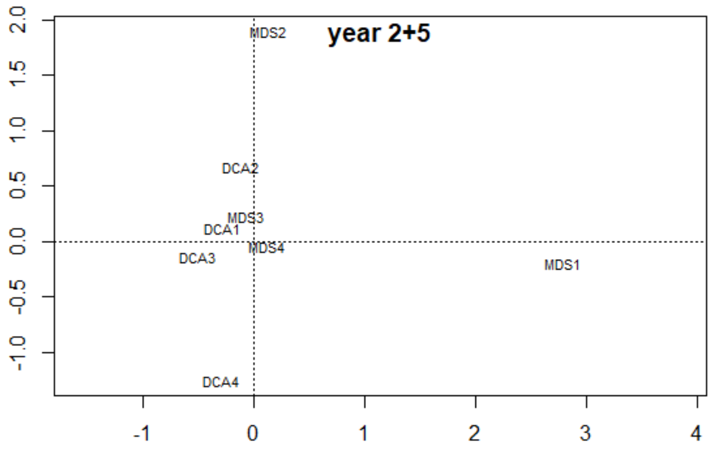


**Figure S4.2.** Principal component analysis (PCA) of ordination axes (gNMDS and DCA).

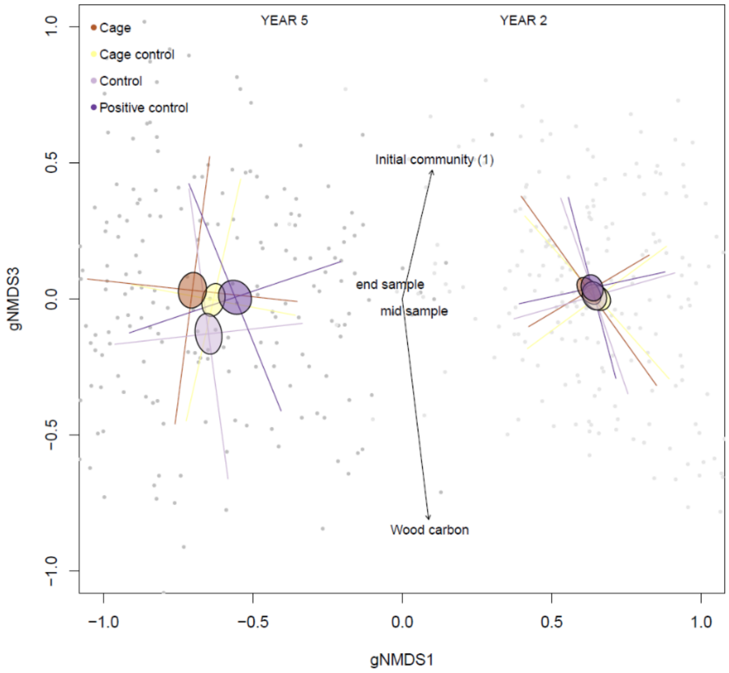


**Figure S4.3.** Ordination biplot of the first and third compositional gradients of fungal OTUs from aspen logs (n = 424) in year 2 and 5 after tree felling (gNMDS of Bray-Curtis dissimilarities; stress = 0.14). Coloured ellipses display the standard error of treatment levels, while error bars display the standard deviation. Vectors and factors are fitted with the 'envfit' function in package VEGAN and are based on a forward selection of linear mixed models of each gNMDS axis. Initial community (1), i.e. DCA1 (Table S1), is multiplied by -1 for visual purposes. Light grey dots represent plot scores of year 2 and dark grey of year 5.

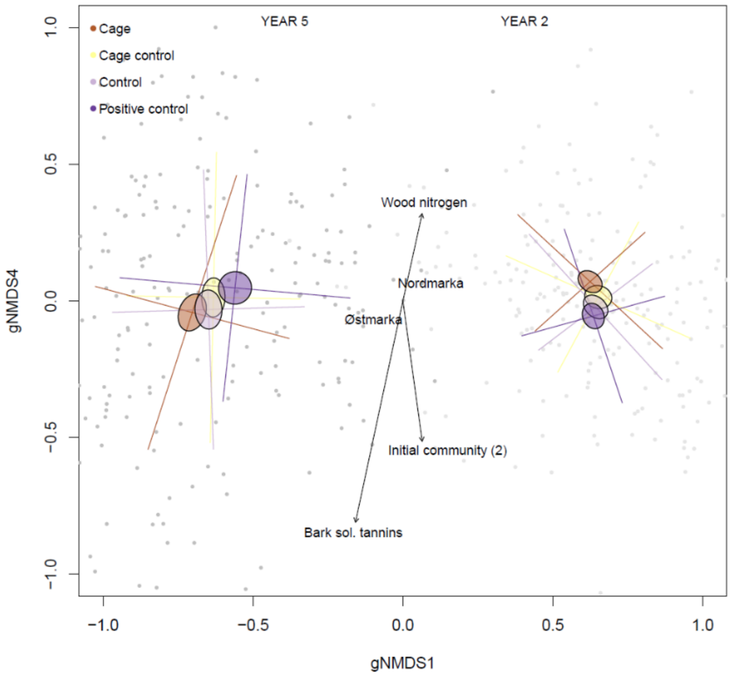


**Figure S4.4.** Ordination biplot of the first and fourth compositional gradients of fungal OTUs from aspen logs (n = 424) in year 2 and 5 after tree felling (gNMDS of Bray-Curtis dissimilarities; stress = 0.14). Coloured ellipses display the standard error of treatment levels, while error bars display the standard deviation. Vectors and factors are fitted with the 'envfit' function in package VEGAN and are based on a forward selection of linear mixed models of each gNMDS axis. Light grey dots represent plot scores of year 2 and dark grey of year 5.

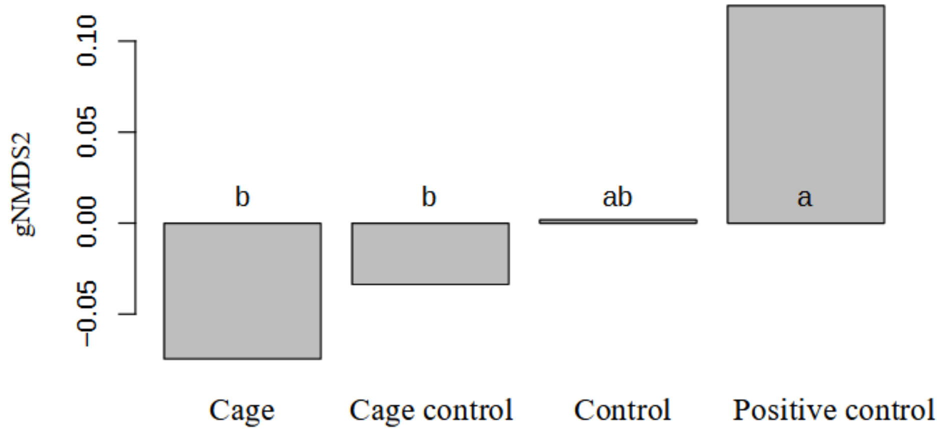


**Figure S4.5**. Tukey HSD test of invertebrate exclusion's effect on gNMDS2 describing variation in community composition of fungal OTUs in aspen wood.

**Table S4.2.** Linear mixed models explaining variation in four compositional gradients (gNMDS; k=4, stress = 1.36) of fungal OTUs from aspen logs in year 2 and 5 after felling. Models are chosen with forward selection (AICc) and Site as random effect. All continuous variables are transformed to reduce skewness and standardised (0 to 1 scale). Reference levels in intercept are: Year (2), Landscape (Østmarka) and Log section (end). Statistic: t. Random effects variance and intraclass correlation coefficient (ICC) is shown.

| | gNMDS1 | | | gNMDS2 | | | gNMDS3 | | | gNMDS4 | | |
|---|---|---|---|---|---|---|---|---|---|---|---|---|
| *Predictors* | *Effect size (95% conf.int.)* | *Statistic* | *p* | *Effect size (95% conf.int.)* | *Statistic* | *p* | *Effect size (95% conf.int.)* | *Statistic* | *p* | *Effect size (95% conf.int.)* | *Statistic* | *p* |
| Intercept | 0.33 (0.21 – 0.46) | 5.22 | **<0.001** | -0.22 (-0.52 – 0.08) | -1.42 | 0.155 | 0.52 (0.28 – 0.76) | 4.26 | **<0.001** | 0.16 (0.03 – 0.29) | 2.49 | **0.013** |
| Year (5) | -1.28 (-1.33 – -1.23) | -47.54 | **<0.001** | | | | | | | | | |
| Landscape (Nordmarka) | 0.23 (0.14 – 0.33) | 4.87 | **<0.001** | | | | | | | 0.15 (0.05 – 0.24) | 3.04 | **0.002** |
| Log section (mid) | 0.08 (0.03 – 0.14) | 3.12 | **0.002** | | | | | | | | | |
| Initial fungal community (1) | | | | | | | -0.26 (-0.45 – -0.08) | -2.77 | **0.006** | | | |
| Initial fungal community (2) | | | | 0.32 (0.06 – 0.58) | 2.44 | **0.015** | | | | -0.45 (-0.67 – -0.24) | -4.09 | **<0.001** |
| Wood carbon | | | | 0.65 (0.22 – 1.08) | 2.94 | **0.003** | -0.69 (-1.04 – -0.34) | -3.88 | **<0.001** | | | |
| Wood nitrogen | | | | | | | | | | 0.21 (0.08 – 0.34) | 3.22 | **0.001** |
| Bark flavonoids | | | | -0.51 (-0.73 – -0.29) | -4.53 | **<0.001** | | | | | | |
| Bark sol. tannins | | | | | | | | | | -0.22 (-0.38 – -0.07) | -2.79 | **0.005** |
| Wood phenolic acids | 0.30 (0.11 – 0.48) | 3.18 | **0.001** | | | | | | | | | |
| **Random Effects** | | | | | | | | | | | | |
| $\sigma^2$ | 0.08 | | | 0.24 | | | 0.16 | | | 0.15 | | |
| $\tau_{00}$ | 0.01 Site | | | 0.02 Site | | | 0.01 Site | | | 0.00 Site | | |
| ICC | 0.12 | | | 0.09 | | | 0.07 | | | 0.03 | | |
| Observations | 424 | | | 424 | | | 424 | | | 424 | | |
| Marginal $R^2$ / Conditional $R^2$ | 0.829 / 0.850 | | | 0.084 / 0.169 | | | 0.052 / 0.122 | | | 0.102 / 0.126 | | |

#### 4.2 Partial constrained ordination

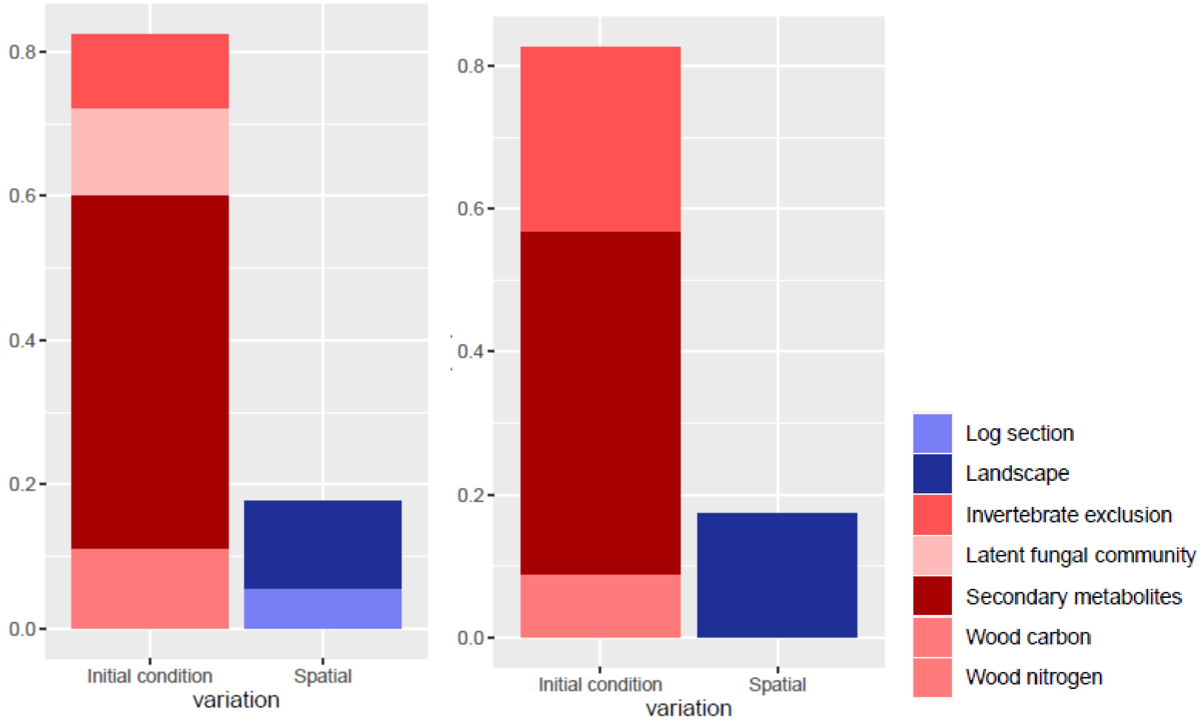


**Figure S4.6.** Proportion of explained variation of fungal community composition in aspen dead wood in year 2 (left) and year 5 (right). The variables included are after a forward model selection approach with constrained ordination (CCA). The variation partitioning is based on inertia units for each variable independent of the other variables in the model.

**Table S4.3.** Variation partitioning with constrained ordination (CCA) of fungal community composition in year 2 and 5 (n=426). Variables are chosen from a forward model selection with *p* as selection criterion. Inertia of constrained variables are modelled alone without any conditioning variables to show variation explained independent of other variables.

| **Variable** | **Inertia** | **Prop. explained** |
|---|---|---|
| Year | 0.5981 | 0.0278 |
| Invertebrate exclusion | 0.1824 | 0.0085 |
| Landscape | 0.1411 | 0.0065 |
| Bark phenolic acids | 0.1060 | 0.0049 |
| Bark flavonoids | 0.1052 | 0.0049 |
| Wood phenolic acids | 0.0967 | 0.0045 |
| Wood flavonoids | 0.0877 | 0.0041 |
| Initial fungal community (1) | 0.0849 | 0.0039 |
| Initial fungal community (2) | 0.0768 | 0.0036 |

| | | |
|---|---|---|
| Wood nitrogen | 0.0651 | 0.0030 |
| Log section | 0.0595 | 0.0028 |
| Unconstrained | 20.0098 | 0.8500 |
| Total | 21.5547 | - * |

*Inertia of constrained and unconstrained axes does not add up to the total inertia because 0.27% of the variation is shared between one or several variables.

##### 4.3 'Envfit'

**Table S4.4**. Environmental variables fitted onto a two-dimensional gNMDS configuration (axis 1 and 2) describing community composition in fungal OTUs. Continuous variables are fitted as vectors with maximum correlation to the configuration. Factor variables are fitted as averages of ordination scores for each level. gNMDS1 and gNMDS2 show the coordinates of each variable. $R^2$ is the correlation coefficient for each variable of the two dimensions. Correlations in bold are significant ($\alpha = 0.05$) from a permutation test (perm. = 999).

| | **gNMDS1** | **gNMDS2** | **$R^2$** |
|---|---|---|---|
| Initial fungal community (1) | -0.95025 | 0.31150 | 0.0012 |
| Initial fungal community (2) | 0.63209 | 0.77489 | 0.0017 |
| Wood salicylates | -0.18034 | 0.98360 | **0.0157** |
| Wood phenolic acids | 0.03789 | 0.99928 | 0.0025 |
| Wood flavonoids | -0.00449 | 0.99999 | 0.0010 |
| Bark phenolic acids | -0.77546 | -0.63139 | 0.0034 |
| Bark flavonoids | -0.12210 | -0.99252 | **0.0481** |
| Wood nitrogen | 0.18781 | 0.98221 | **0.0146** |
| Wood carbon | 0.11768 | 0.99305 | **0.0372** |
| Bark MeOH-soluble condensed tannins | -0.60953 | 0.79276 | 0.0106 |
| Bark MeOH-insoluble condensed tannins | -0.44319 | 0.89643 | 0.0027 |
| Landscape (Østmarka) | -0.0982 | 0.0850 | |
| (Nordmarka) | 0.0928 | -0.0803 | **0.0201** |
| Invertebrate exclusion (cage) | -0.0391 | -0.0749 | |
| (Cage control) | 0.0097 | -0.0330 | |
| (Control) | -0.0029 | 0.0086 | |
| (Positive control) | 0.0376 | 0.1151 | 0.0071 |
| Section (end) | -0.0407 | 0.0126 | |

| | | | |
|---|---|---|---|
| (mid) | 0.0404 | -0.0125 | 0.0023 |

**Table S4.5**. Environmental variables fitted onto a two-dimensional gNMDS configuration (axis 1 and 3) describing community composition in fungal OTUs. Continuous variables are fitted as vectors with maximum correlation to the configuration. Factor variables are fitted as averages of ordination scores for each level. gNMDS1 and gNMDS3 show the coordinates of each variable. $R^2$ is the correlation coefficient for each variable of the two dimensions. Correlations in bold are significant (α = 0.05) from a permutation test (perm. = 999).

| | gNMDS1 | gNMDS3 | r2 |
|---|---|---|---|
| Initial fungal community (1) | -0.20736 | -0.97827 | 0.0099 |
| Initial fungal community (2) | 0.39840 | 0.91721 | 0.0027 |
| Wood salicylates | -0.52450 | 0.85141 | 0.0017 |
| Wood phenolic acids | 0.02746 | -0.99962 | 0.0030 |
| Wood flavonoids | -0.00475 | -0.99999 | 0.0006 |
| Bark phenolic acids | -0.32383 | -0.94611 | 0.0099 |
| Bark flavonoids | -0.41384 | 0.91035 | 0.0034 |
| Wood nitrogen | 0.23212 | 0.97269 | 0.0064 |
| Wood carbon | 0.10805 | -0.99415 | **0.0280** |
| Bark MeOH-soluble condensed tannins | -0.31077 | 0.95048 | **0.0234** |
| Bark MeOH-insoluble condensed tannins | -0.27393 | 0.96175 | 0.0044 |
| Landscape (Østmarka) | -0.0982 | 0.0105 | |
| (Nordmarka) | 0.0928 | -0.0100 | **0.0134** |
| Invertebrate exclusion (cage) | -0.0391 | 0.0309 | |
| (Cage control) | 0.0097 | 0.0011 | |
| (Control) | -0.0029 | -0.0583 | |
| (Positive control) | 0.0376 | 0.0235 | 0.0029 |
| Section (end) | -0.0407 | 0.0483 | |
| (mid) | 0.0404 | -0.0478 | 0.0057 |

**Table S4.6**. Environmental variables fitted onto a two-dimensional gNMDS configuration (axis 1 and 4) describing community composition in fungal OTUs. Continuous variables are fitted as vectors with maximum correlation to the configuration. Factor variables are fitted as averages of ordination scores for each level. gNMDS1 and gNMDS4 show the coordinates of each variable. $R^2$ is the correlation coefficient for each variable of the two dimensions. Correlations in bold are significant (α = 0.05) from a permutation test (perm. = 999).

| | gNMDS1 | gNMDS4 | r2 |
|---|---|---|---|
| Initial fungal community (1) | -0.42569 | -0.90487 | 0.0028 |
| Initial fungal community (2) | 0.12454 | -0.99221 | **0.0211** |
| Wood salicylates | -0.12206 | -0.99252 | **0.0207** |
| Wood phenolic acids | 0.15889 | 0.98730 | 0.0001 |
| Wood flavonoids | -0.00182 | -1.00000 | 0.0037 |
| Bark phenolic acids | -0.24575 | -0.96933 | **0.0154** |
| Bark flavonoids | -0.21631 | -0.97633 | 0.0100 |
| Wood nitrogen | 0.19719 | 0.98036 | 0.0083 |
| Wood carbon | 0.51398 | 0.85780 | 0.0018 |
| Bark MeOH-soluble condensed tannins | -0.19171 | -0.98145 | **0.0535** |
| Bark MeOH-insoluble condensed tannins | -0.17609 | -0.98437 | 0.0095 |
| Landscape (Østmarka) | -0.0982 | -0.0691 | |
| (Nordmarka) | 0.0928 | 0.0653 | **0.0200** |
| Invertebrate exclusion (cage) | -0.0391 | 0.0141 | |
| (Cage control) | 0.0097 | 0.0134 | |
| (Control) | -0.0029 | -0.0267 | |
| (Positive control) | 0.0376 | -0.0037 | 0.0015 |
| Section (end) | -0.0407 | -0.0006 | |
| (mid) | 0.0404 | 0.0006 | 0.0024 |

#### 5. Model diagnostics

We examined all linear mixed models (LMMs) of wood density and fungal OTU richness for violations of model assumptions (Zuur et al. 2010). We checked for multicollinearity with generalized variance inflation factors. We examined serial plots of the residuals to look for non-independence. We examined residual plots, q-q plots and residual distributions to check for deviations to the assumptions of homoscedasticity and linearity of model residuals, random effects and fixed effects. If several model assumptions seemed to be violated, we also tested for model fit by posterior predictive simulations as described in Bates et al. (2014).

In general, model assumptions were met, except for extreme deviations of the assumed linearity of fixed effect residuals for all variables (Figure S5.1). This was not surprising as the continuous variables were derived from fewer samples than those for wood density and fungal OTUs (Figure S1). Bark and wood variables had 17 and 50 unique values, respectively (Table S3), which meant that the values would be non-randomly duplicated before fitting them into the models.

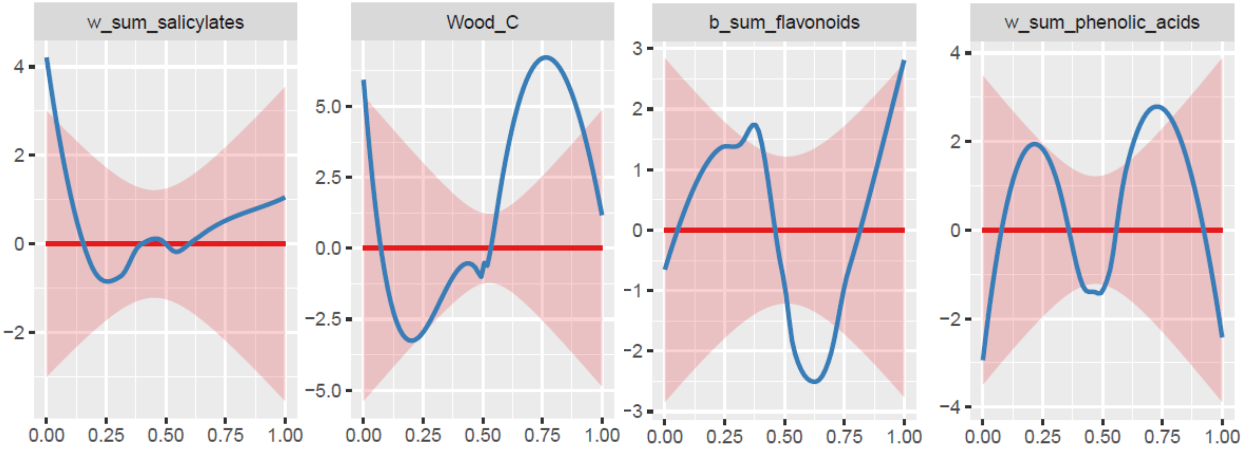


**Figure S5.1.** Extreme deviations from the assumptions of linearity in linear mixed models. Here is an example showing residuals of continuous fixed effects from a model explaining variation in fungal OTU richness in year 5 after tree felling.

To evaluate the certainty of the models explaining variation in wood density and fungal OTU richness, we therefore ran new models: two without random effects ('unaggregated') and two with aggregated datasets (using the mean of the response variables), investigating wood and bark variables separately for each response variable. Then, we compared effect sizes and *p* values of continuous explanatory variables of the new and the original models.

##### 5.1 Fungal OTU richness

New models for explaining fungal OTU richness in year 2 and year 5 were made for both bark and wood variables.

In year 2, most effect sizes were similar between models using the original ('unaggregated with random term'), unaggregated and aggregated datasets. Initial fungal community (1), wood flavonoids and bark

MeOH-soluble condensed tannins had divergent results between models of the three datasets (Figure S5.2). However, the only significant continuous variable explaining fungal OTU richness in year 2, wood carbon, had similar effect sizes and was also significant without random term and in an aggregated dataset (Figure S5.2).

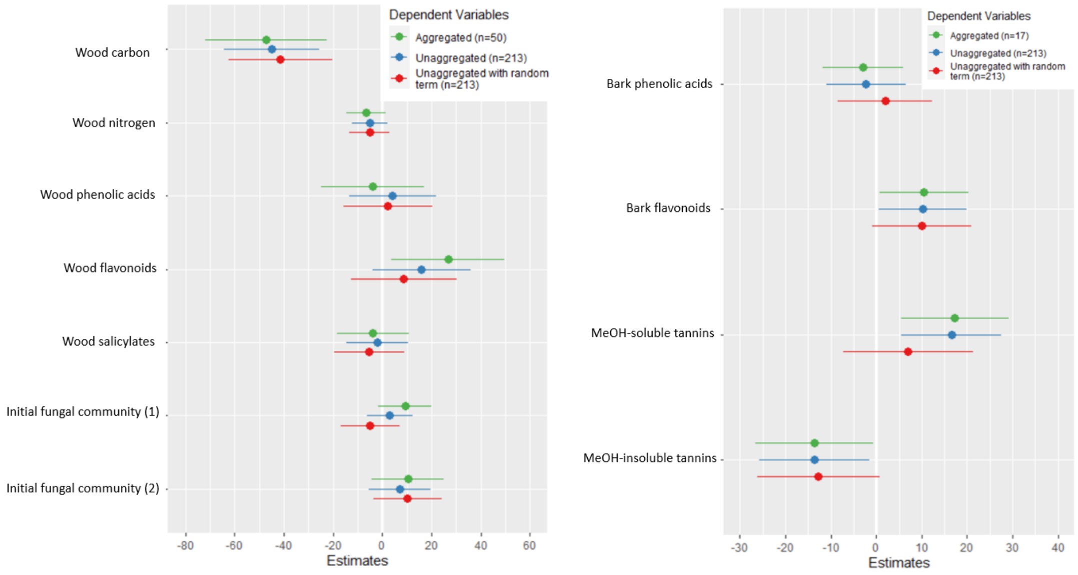


**Figure S5.2.** Comparing effect sizes of wood (left) and bark (right) variables in three different linear models explaining fungal OTU richness in year 2: the original model (LMM) used in the article ('unaggregated with random term'), the same model without random term ('unaggregated') and a model where the dataset was aggregated by the number of unique values of the explanatory variables ('aggregated'). Aggregation was done seperately to test wood and bark variables because they had different number of unique values (50 and 17, respectively).

In year 5, the models were in agreement with the direction of each variable's effect (positive or negative), but not in *p* values (Figure S5.3). Notably, wood phenolic acids was not significant in the aggregated dataset as the model estimate was similar, but the variation was larger.

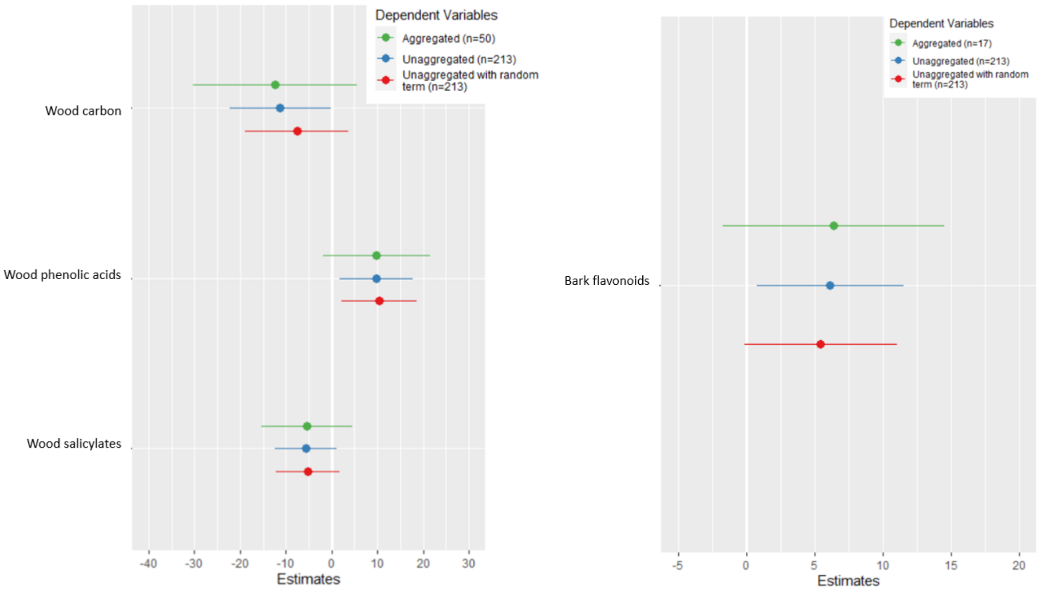

**Figure S5.3.** Comparing effect sizes of wood (left) and bark (right) variables in three different linear models explaining fungal OTU richness in year 5: the original model (LMM) used in the article ('unaggregated with random term'), the same model without random term ('unaggregated') and a model where the dataset was aggregated by the number of unique values of the explanatory variables ('aggregated'). Aggregation was done seperately to test wood and bark variables because they had different number of unique values (50 and 17, respectively).

#### 5.1 Wood density

New models for explaining wood density in year 2 and year 5 were made for both bark variables (as no wood variables were included in the model selection procedure).

In year 2, the variation was much larger in aggregated datasets, which resulted in no significant p values (Table S5.1). However, the aggregated dataset had undergone a strong transformation of the response variable (mean wood density per tree individual), which reduced the sample size to only 17 and removes potentially important information on the ecological variation and response's error distribution (Reitan and Nielsen 2016). Therefore, we decided to investigate the wood density models further with the original dataset fitted with polynomial splines (James et al. 2013). We used function 'bs' and increased degrees of freedom until assumption of linearity in fixed effect residuals were met (Team 2021).

**Table S5.1.** Comparing effect sizes bark variables in three different linear models explaining wood density in year 2: the original model (LMM) used in the article ('unaggregated with random term'), the same model without random term ('unaggregated') and a model where the dataset was aggregated by the number of unique bark values ('aggregated').

| | Aggregated (n=17) | | | Unaggregated (n=213) | | | Unaggregated with random term (n=213) | | |
|---|---|---|---|---|---|---|---|---|---|
| *Predictors* | *Effect size (95% conf.int.)* | *Statistic* | *p* | *Effect size (95% conf.int.)* | *Statistic* | *p* | *Effect size (95% conf.int.)* | *Statistic* | *p* |
| Intercept | 0.37 (0.34 – 0.40) | 24.32 | **<0.001** | 0.37 (0.36 – 0.38) | 72.53 | **<0.001** | 0.36 (0.35 – 0.37) | 65.75 | **<0.001** |
| Bark phenolic acids | 0.02 (-0.03 – 0.07) | 0.92 | 0.375 | 0.03 (0.01 – 0.04) | 3.86 | **<0.001** | 0.02 (0.01 – 0.04) | 3.45 | **0.001** |
| Bark insol. tannins | 0.03 (-0.03 – 0.08) | 1.08 | 0.298 | 0.03 (0.01 – 0.04) | 3.16 | **0.002** | 0.04 (0.02 – 0.05) | 5.12 | **<0.001** |

Both bark phenolic acids and MeOH-insoluble condensed tannins were fitted with the 7th order splines in a model that was a large improvement from the original model (ΔAIC = 31.9). As in the original model, tannins showed a significant positive relationship with wood density at the 1st, 2nd and 7th degrees (Table S5.2). Wood phenolic acids, however, had a more complicated relationship with wood density, predicting both positive and negative relationships at different values (Figure S5.4, Table S5.2). We conclude that the positive effect of bark phenolic acids on wood density in year 2 should be interpreted with caution.

**Table S5.2.** Polynomial spline regression explaining variation in wood density in year 2. Fitted with site as random effect and the 7th order splines of bark MeOH-insoluble condensed tannins and bark phenolic acids. Significant t values in bold (α = 0.05).

| Predictor | Estimate | Std. error | DF | t value |
|---|---|---|---|---|
| (Intercept) | 0.358904 | 0.018288 | 134.433661 | **19.626** |
| Bark MeOH-soluble tannins (1) | -0.071702 | 0.052262 | 161.759581 | -1.372 |
| Bark MeOH-soluble tannins (2) | 0.129107 | 0.050634 | 152.391323 | **2.550** |

| | | | | |
|---|---|---|---|---|
| Bark MeOH-soluble tannins (3) | 0.059815 | 0.011702 | 109.693184 | **5.112** |
| Bark MeOH-soluble tannins (4) | 0.050483 | 0.063465 | 165.706579 | 0.795 |
| Bark MeOH-soluble tannins (5) | 0.019033 | 0.067867 | 163.370351 | 0.280 |
| Bark MeOH-soluble tannins (6) | 0.031826 | 0.027710 | 141.920586 | 1.149 |
| Bark MeOH-soluble tannins (7) | 0.091725 | 0.021830 | 172.682948 | **4.202** |
| Bark phenolic acids (1) | -0.047372 | 0.014001 | 134.988207 | **-3.383** |
| Bark phenolic acids (2) | -0.080764 | 0.028181 | 148.094988 | **-2.866** |
| Bark phenolic acids (3) | 0.077193 | 0.037613 | 134.148856 | **2.052** |
| Bark phenolic acids (4) | -0.096178 | 0.015490 | 125.326854 | **-6.209** |
| Bark phenolic acids (5) | 0.134906 | 0.037663 | 135.172828 | **3.582** |
| Bark phenolic acids (6) | -0.275165 | 0.100356 | 162.183855 | **-2.742** |
| Bark phenolic acids (7) | 0.008413 | 0.011926 | 136.454474 | 0.705 |

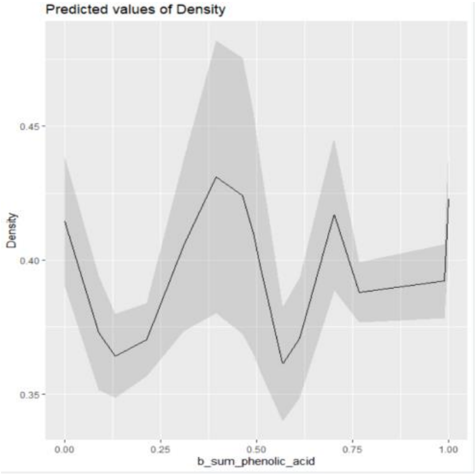


**Figure S5.4.** Expected wood density in year 2 at different levels of bark phenolic acids, estimated with a 7th ordered spline regression with site as random effect.

In year 5, bark MeOH-insoluble condensed tannins had very similar effect sizes between aggregated, unaggregated and the original models (Table S5.3). The positive effect that tannins had on wood density was strong, but not significant ($p = 0.071$) and we decided to compare the model with spline regression too. The tannins variable was fitted with the 4th order splines and improved model fit ($\Delta AIC = 4.2$) and did indeed predict positive relationships between wood density at both the 2nd and 4th degrees (Table S5.4).

**Table S5.3.** Comparing effect sizes of bark MeOH-soluble condensed tannins in three different linear models explaining wood density in year 5: the original model (LMM) used in the article (‘unaggregated with random term’), the same model without random term (‘unaggregated’) and a model where the dataset was aggregated by the number of unique values of tannins (‘aggregated’).

| Predictors | Aggregated (n=17) Effect size (95% conf.int.) | Statistic | *p* | Unaggregated (n=213) Effect size (95% conf.int.) | Statistic | *p* | Unaggregated with random term (n=213) Effect size (95% conf.int.) | Statistic | *p* |
|---|---|---|---|---|---|---|---|---|---|
| Intercept | 0.34 (0.31 – 0.37) | 23.94 | **<0.001** | 0.33 (0.32 – 0.35) | 48.21 | **<0.001** | 0.33 (0.31 – 0.34) | 43.33 | **<0.001** |
| Bark insol. tannins | 0.04 (-0.00 – 0.09) | 1.94 | 0.071 | 0.05 (0.03 – 0.07) | 4.40 | **<0.001** | 0.06 (0.04 – 0.08) | 6.07 | **<0.001** |

**Table S5.3**. Polynomial spline regression explaining variation in wood density in year 5. Fitted with site as random effect and the 4th order splines of bark MeOH-insoluble condensed tannins. Significant values in bold (α = 0.05)

| Predictors | Estimate | Std. error | DF | t value |
|---|---|---|---|---|
| (Intercept) | 0.31271 | 0.01055 | 178.35400 | **29.652** |
| Bark MeOH-soluble tannins (1) | 0.03871 | 0.02717 | 205.07550 | 1.425 |
| Bark MeOH-soluble tannins (2) | 0.09161 | 0.02376 | 198.55914 | **3.856** |
| Bark MeOH-soluble tannins (3) | -0.02194 | 0.02383 | 202.47996 | -0.921 |
| Bark MeOH-soluble tannins (4) | 0.09910 | 0.01187 | 193.33802 | **8.352** |

#### 6. Model of fungal OTU richness

**Table S6.** Variation in fungal OTU richness in aspen wood year 2 (left) and year 5 (right) after tree felling. Estimated from a linear mixed model with site as random effect.

| | Year 2 | | Year 5 | |
|---|---|---|---|---|
| *Predictors* | *Effect size (95% conf. int.)* | *p* | *Effect size (95% conf. int.)* | *p* |
| Intercept | 92.49 (71.33 – 113.65) | **<0.001** | 47.54 (39.21 – 55.87) | **<0.001** |
| Treatment (cage control) | 0.06 (-5.43 – 5.54) | 0.984 | 2.23 (-1.23 – 5.68) | 0.206 |
| Treatment (control) | -1.30 (-6.79 – 4.20) | 0.644 | 1.56 (-1.96 – 5.09) | 0.384 |
| Treatment (OH control) | -2.23 (-7.73 – 3.27) | 0.426 | 1.22 (-2.33 – 4.77) | 0.500 |
| Log section (mid) | 4.70 (0.91 – 8.50) | **0.015** | | |
| Landscape (Nordmarka) | -7.87 (-15.27 – -0.47) | **0.037** | | |
| Initial fungal community (1) | -4.93 (-16.82 – 6.95) | 0.416 | | |
| Initial fungal community (2) | 10.31 (-3.60 – 24.22) | 0.146 | | |
| Wood carbon | -41.44 (-62.47 – -20.41) | **<0.001** | -7.55 (-19.00 – 3.90) | 0.196 |
| Wood nitrogen | -5.25 (-13.45 – 2.95) | 0.209 | | |
| Wood phenolic acids | 2.32 (-15.82 – 20.46) | 0.802 | 10.63 (2.27 – 18.98) | **0.013** |
| Wood salicylates | -5.34 (-19.67 – 8.99) | 0.465 | -5.48 (-12.48 – 1.51) | 0.125 |
| Wood flavonoids | 8.70 (-12.83 – 30.23) | 0.428 | | |
| Bark phenolic acids | 1.96 (-8.45 – 12.37) | 0.712 | | |
| Bark flavonoids | 10.04 (-0.85 – 20.92) | 0.071 | 5.24 (-0.39 – 10.87) | 0.068 |
| Bark sol. tannins | 7.03 (-7.20 – 21.26) | 0.333 | | |
| Bark insol. tannins | -12.73 (-26.15 – 0.68) | 0.063 | | |
| **Random Effects** | | | | |
| $\sigma^2$ | 196.52 | | 85.81 | |
| $\tau_{00}$ | 52.61 $_{Site}$ | | 5.39 $_{Site}$ | |
| ICC | 0.21 | | 0.06 | |
| N | 30 $_{Site}$ | | 30 $_{Site}$ | |
| Observations | 213 | | 213 | |
| Marginal $R^2$ / Conditional $R^2$ | 0.207 / 0.375 | | 0.062 / 0.118 | |
